# A Spiking Neuron and Population Model based on the Growth Transform Dynamical System

**DOI:** 10.1101/523944

**Authors:** Ahana Gangopadhyay, Darshit Mehta, Shantanu Chakrabartty

**Affiliations:** Department of Electrical and Systems Engineering, Washington University in St. Louis; Department of Biomedical Engineering, Washington University in St. Louis

**Keywords:** Spiking neuron model, Growth Transforms, energy-minimization, dynamical system, network model, neural dynamics, associative memory, adaptation

## Abstract

In neuromorphic engineering, neural populations are generally modeled in a bottom-up manner, where individual neuron models are connected through synapses to form large-scale spiking networks. Alternatively, a top-down approach treats the process of spike generation and neural representation of excitation in the context of minimizing some measure of network energy. However, these approaches usually define the energy functional in terms of some statistical measure of spiking activity (ex. firing rates), which does not allow independent control and optimization of neurodynamical parameters. In this paper, we introduce a new spiking neuron and population model where the dynamical and spiking responses of neurons can be derived directly from a network objective or energy functional of continuous-valued neural variables like the membrane potential. The key advantage of the model is that it allows for independent control over three neuro-dynamical properties: (a) control over the steady-state population dynamics that encodes the minimum of an exact network energy functional; (b) control over the shape of the action potentials generated by individual neurons in the network without affecting the network minimum; and (c) control over spiking statistics and transient population dynamics without affecting the network minimum or the shape of action potentials. At the core of the proposed model are different variants of Growth Transform dynamical systems that produce stable and interpretable population dynamics, irrespective of the network size and the type of neuronal connectivity (inhibitory or excitatory). In this paper, we present several examples where the proposed model has been configured to produce different types of single-neuron dynamics as well as population dynamics. In one such example, the network is shown to adapt such that it encodes the steady-state solution using a reduced number of spikes upon convergence to the optimal solution. In this paper we use this network to construct a spiking associative memory that uses fewer spikes compared to conventional architectures, while maintaining high recall accuracy at high memory loads.

## 1 Introduction

Spiking neural networks that emulate neural ensembles have been studied extensively within the context of dynamical systems [1], and modeled as a set of differential equations that govern the temporal evolution of its state variables. For a single neuron, the state variables are usually its membrane potential and the conductances of ion channels that mediate changes in the membrane potential via flux of ions across the cell membrane. A vast body of literature, ranging from the classical Hodgkin-Huxley model [2], FitzHugh-Nagumo model [3], Izhikevich model [4] to simpler integrate-and-fire models [5], treats the problem of single-cell excitability at various levels of detail and biophysical plausibility. Individual neuron models are then connected through synapses, bottom-up, to form large-scale spiking neural networks.

An alternative to this bottom-up approach is a top-down approach that treats the process of spike generation and neural representation of excitation in the context of minimizing some measure of network energy. The rationale for this approach is that physical processes occurring in nature have a tendency to self-optimize towards a minimum-energy state. This principle has been used to design neuromorphic systems where the state of a neuron in the network is assumed to be either binary in nature (spiking or not spiking) [6], or replaced by its average firing rate [7]. However, in all of these approaches, the energy functionals have been defined with respect to some statistical measure of neural activity, for example spike rates, instead of continuous-valued neuronal variables like the membrane potential. As a result in these models, it is difficult to independently control different neuro-dynamical parameters, for example the shape of the action-potential, bursting activity or adaptation in neural activity, without affecting the network solution.

In [8], we proposed a model of a Growth Transform (GT) neuron which reconciled the bottom-up and top-down approaches such that the dynamical and spiking responses were derived directly from a network objective or an energy functional. Each neuron in the network implements an asynchronous mapping based on polynomial Growth Transforms, which is a fixed-point algorithm for optimizing polynomial functions under linear and/or bound constraints [9, 10]. It was shown in [8] that a network of GT neurons can solve binary classification tasks while producing stable and unique neural dynamics (for example, noise-shaping, spiking and bursting) that could be interpreted using a classification margin. However, in the previous formulation, all of these neuro-dynamical properties were directly encoded into the network energy function. As a result, the formulation did not allow independent control and optimization of different neuro-dynamics. In this paper, we address these limitations by proposing a novel GT spiking neuron and population model, along with a neuromorphic framework, according to the following steps:

- We first remap the synaptic interactions in a standard spiking neural network in a manner that the solution (steady-state attractor) could be encoded as a first-order condition of an optimization problem. We show that this network objective function or energy functional can be interpreted as the total extrinsic power required by the remapped network to operate, and hence a metric to be minimized.
- We then introduce a dynamical system model based on Growth Transforms that evolves the network towards this steady-state attractor under the specified constraints. The use of Growth Transforms ensures that the neuronal states (membrane potentials) involved in the optimization are always bounded and that each step in the evolution is guaranteed to reduce the network energy.
- We then show how gradient discontinuity in the regularization function of the network energy functional can be used to incorporate the shape of the action potential while maintaining the local convexity of the energy functional and hence the location of the steady-state attractor.
- Finally, we use the properties of Growth Transforms to generalize the model to a continuous-time dynamical system. The formulation will then allow for modulating the spiking and the population dynamics across the network without affecting network convergence to the steady-state attractor.

We show that the proposed framework can be used to implement a network of coupled neurons that can exhibit memory, global adaptation, and other interesting population dynamics under different initial conditions and based on different network states. We also illustrate how decoupling transient spiking dynamics from the network solution and spike-shapes could be beneficial by using the model to design a spiking associative memory network that can recall a large number of patterns with high accuracy while using fewer spikes than traditional associative memory networks. This paper is also accompanied by a publicly available software implementing the proposed model [11] using MATLAB^©^. Users can experiment with different inputs and network parameters to explore and create other unique dynamics than what has been reported in this paper. In the future, we envision that the model could be extended to incorporate spike-based learning within an energy-minimization framework similar to the framework used in traditional machine learning models [12]. This could be instrumental in bridging the gap between neuromorphic algorithms and traditional energy-based machine learning models.

## 2 Methods

In this section, we present the network energy functional by remapping the synaptic interactions of a standard spiking neural network and then propose a Growth Transform based dynamical system for minimizing this objective. For the rest of the paper, we will follow the mathematical notations as summarized below.

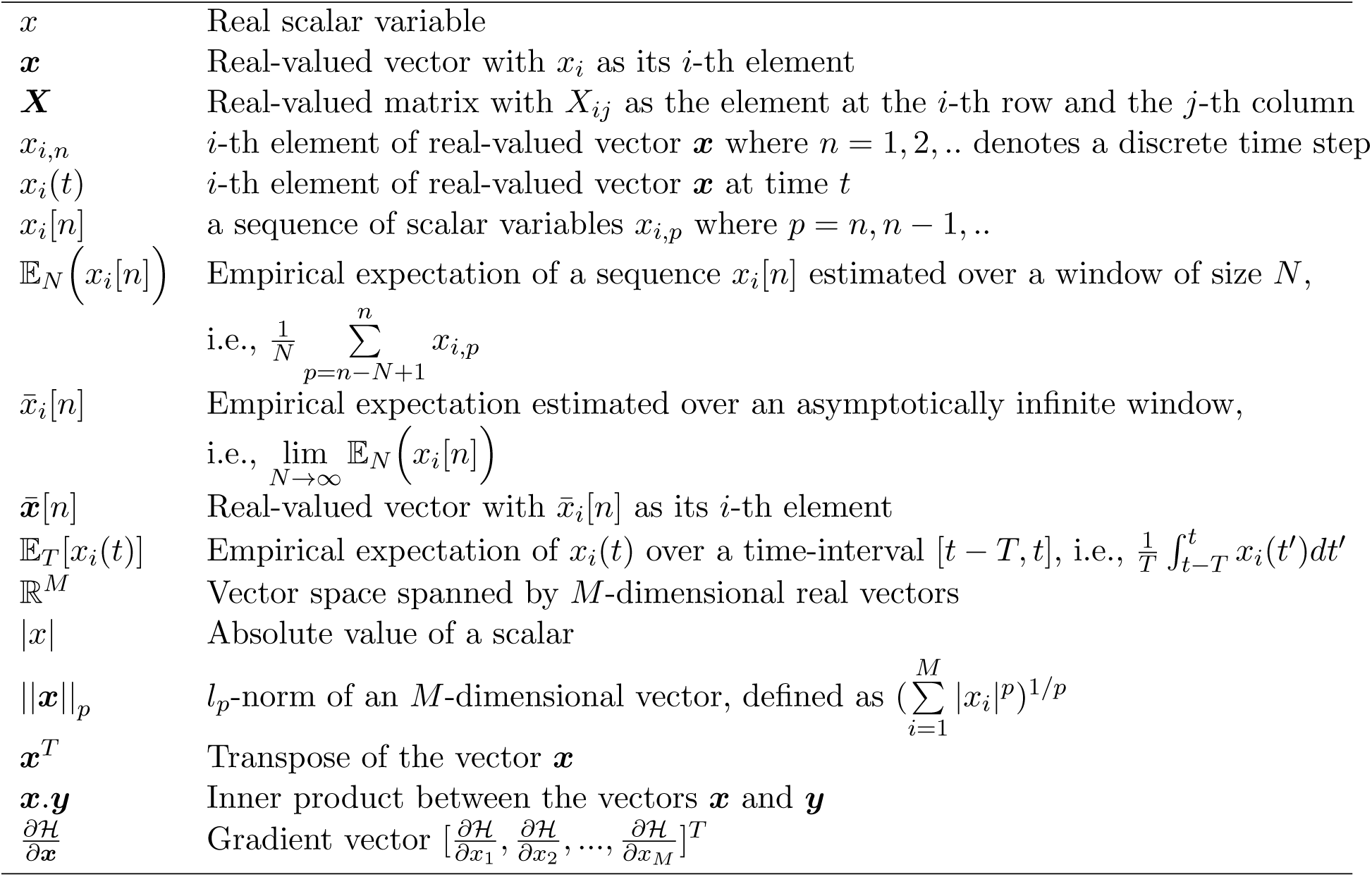
Notations.

### 2.1 Remapping synaptic interactions in a standard spiking network

In generalized threshold models like the Spike Response Model [13], the membrane voltage is given using response kernels that accurately model the post-synaptic responses due to pre-synaptic input spikes, external driving currents and the shape of the spike - the latter term being also used to model refractoriness. However, in simpler adaptations of spiking neuron models, the spike shape is often disregarded, and the membrane potentials are written as simple linear post-synaptic integrations of input spikes and external currents [14, 15]. We consider a similar model where *v*_*i*_ ∈ ℝ represents an inter-cellular membrane potential corresponding to neuron *i* in a network of *M* neurons. The *i*-th neuron receives spikes from the *j*-th neuron that are modulated by a synapse whose strength or weight is denoted by *W*_*ij*_ ∈ R. Assuming that the synaptic weights are constant, the following discrete-time temporal equation governs the dynamics when the membrane potential increases [16, 17]

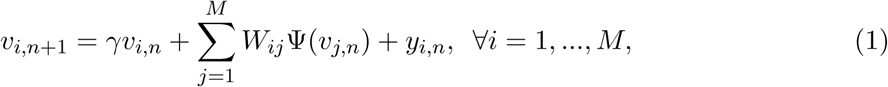

where *v*_*i,n*_ ≡ *v*_*i*_(*n*Δ*t*) and *v*_*i,n*+1_ ≡ *v*_*i*_ ((*n* + 1)Δ*t*), Δ*t* being the time increment between two time-steps. *y*_*i,n*_ represents the depolarization due to an external stimulus that can be viewed as *y*_*i,n*_ = *R*_*mi*_*I*_*i,n*_, where *I*_*i,n*_ ∈ ℝ is the current stimulus at the *n*-th time-step and *R*_*mi*_ ∈ ℝ is the membrane resistance of the *i*-th neuron. Here, 0 *≤ γ ≤* 1 denotes the leakage factor and Ψ(.) denotes a simple spiking function that is positive only when the voltage *v*_*j,n*_ exceeds a threshold and 0 otherwise. Note that in (1), the filter Ψ(.) implicitly depends on the pre-synaptic spike-times through the pre-synaptic membrane voltage *v*_*j,n*_. Such a spiking neural network model is shown in Figure 1(a).

**Figure 1:**
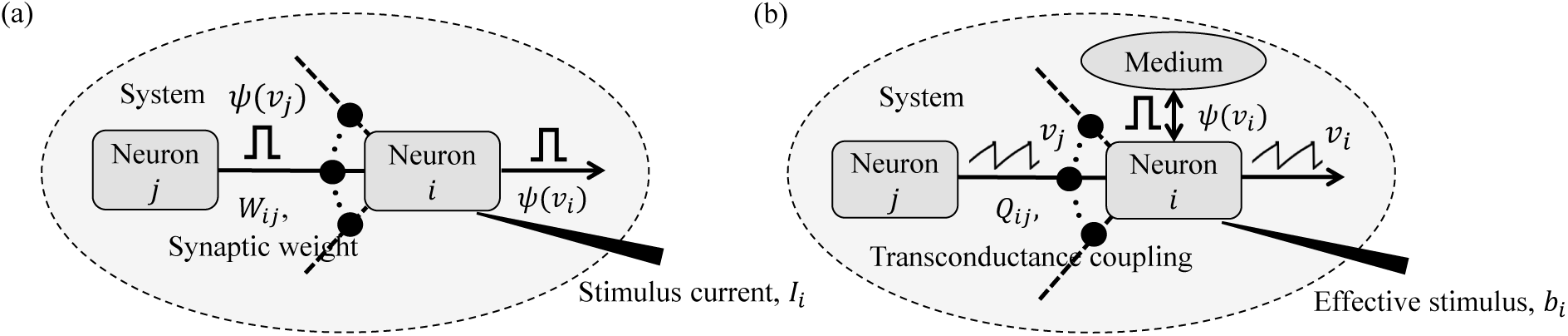
(a) Simple but general model of a spiking neural network; (b) Compartmental network model obtained after remapping.

We further enforce that the membrane potentials are bounded by *v*_*c*_ as

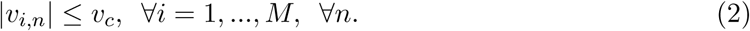

Note that in biological neural networks, the membrane potentials are also bounded [18].

If Ψ(.) was a smooth function of the membrane potential, *v*_*i,n*_ would track the net input at every instant. For a non-smooth Ψ(.), however, we make the additional assumption that the temporal expectation of *v*_*i,n*_ encodes the net input over a sufficiently large time-window. Considering 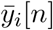 to be the empirical expectation of the external input estimated at the *n*-th time-window, and under the bound constraints outlined in (2), we can get the following relation (justification in Appendix A)

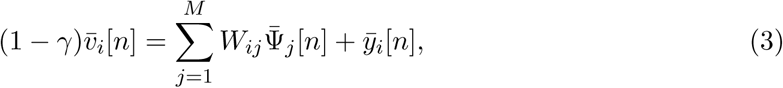

where 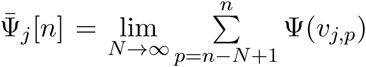. To reduce notational clutter, we will re-write (3) in a matrix form as

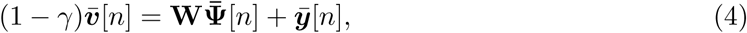

where 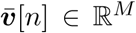 is the vector of mean membrane potentials for a network of *M* neurons, **W** ∈ ℝ^*M*^ *×* ℝ^*M*^ is the synaptic weight matrix for the network, 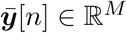 is the vector of mean external inputs for the *n*-th time-window and 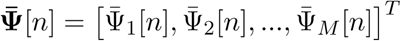 is the vector of mean spike currents. As Ψ(.) is a non-linear function of the membrane potential, it is difficult to derive an exact network energy functional corresponding to (4). However, if we assume that the synaptic weight matrix ***W*** is invertible, we can re-write Eq. (4) as

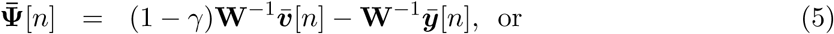

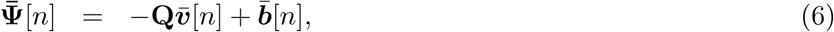

where **Q** = −(1 − *γ*)**W**^−1^, and 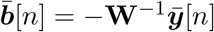 is the effective external current stimulus. Note that in case ***W*** is not invertible, ***W***^−1^ could represent a pseudo-inverse. For the *i*-th neuron, (6) is equivalent to

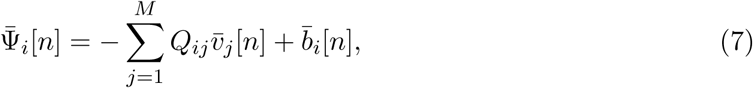

subject to the bound constraint |*v*_*i,n*_| *≤ v*_*c*_ ∀*i, n*. In the subsequent sections, we show that (7) can be viewed as the first-order condition of the following network objective function or energy functional

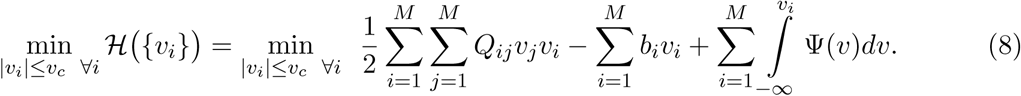

The network energy functional *ℋ*(.) in (8) also admits a physical interpretation, as shown in Figure 1(b). Each neuron *i* receives a voltage input from the neuron *j* through a synapse that can be modeled by a transconductance *Q*_*ij*_ [19]. The neuron *i* also receives an electrical current stimulus *b*_*i*_ and exchanges a voltage-dependent ionic-current with its medium, denoted by Ψ(*v*_*i*_). Then, the function *ℋ* (.) in Eq. (8) represents the extrinsic (or metabolic) power supplied to the network, comprising the following three components: (a) Power dissipation due to coupling between neurons; (b) Power injected to or extracted from the system as a result of external stimulation; and (c) Power dissipated due to neural responses.

### 2.2 Neuron model using the Growth Transform dynamical system

In order to solve the energy minimization problem given in (8) under the constraints given in (2), we first propose a dynamical system based on polynomial Growth Transforms. We also show how the dynamical system evolves over time to satisfy (7) as a first-order condition.

Growth Transforms are multiplicative updates derived from the well-known Baum-Eagon inequality [9, 20] that optimize a Lipschitz continuous cost function under linear and/or bound constraints on the optimization variables. Each neuron in the network implements a continuous mapping based on Growth Transforms, ensuring that the network evolves over time to reach an optimal solution of the energy functional within the constraint manifold. The summary of the proposed model is presented in Table 1 and the detailed derivation is provided in Appendix B.

**Table 1:**
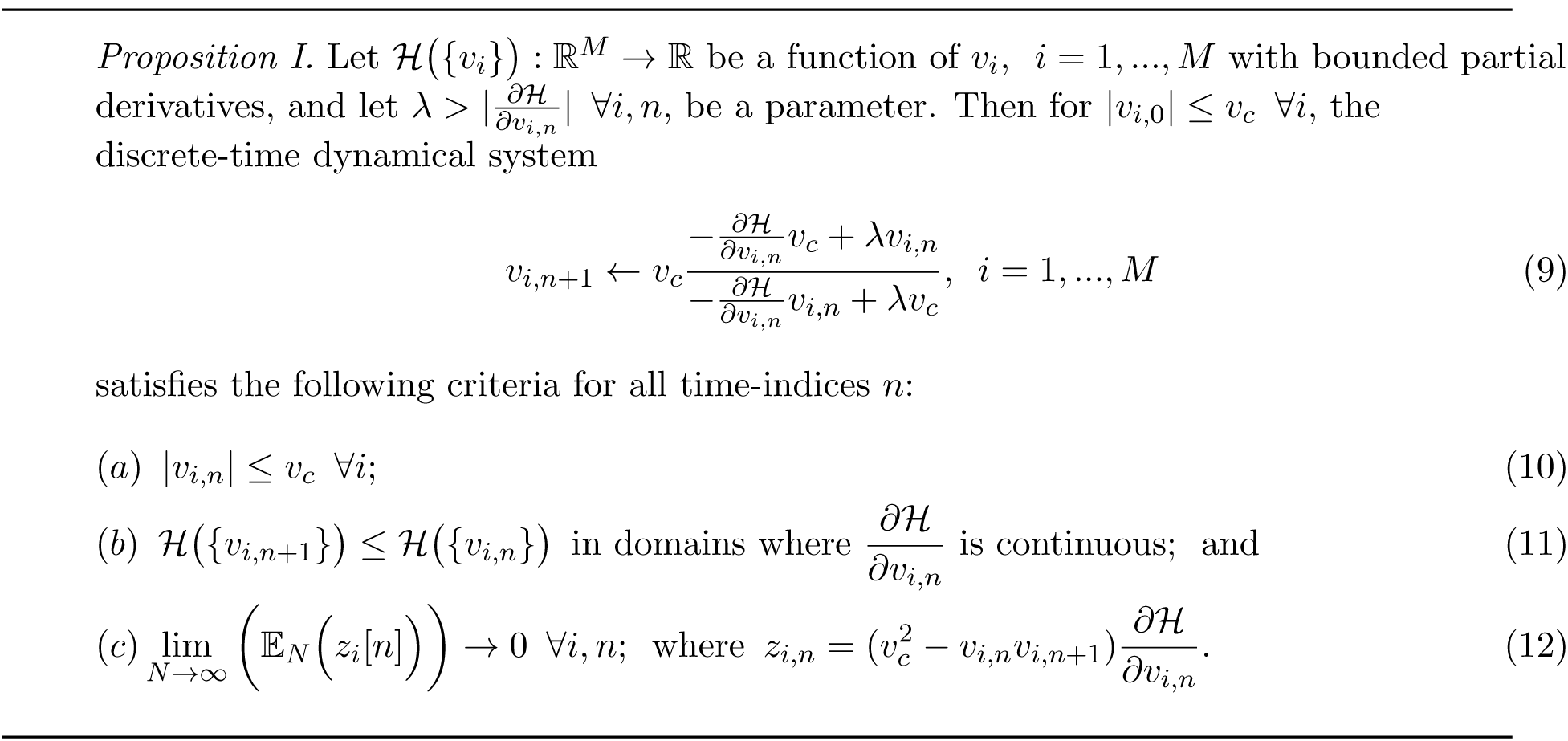
Discrete-time Growth Transform dynamical system (Proof in Appendix B)

#### 2.2.1 Growth Transform spiking neuron model

Considering the *n*-th iteration of the update equation in (9) as the *n*-th time-step for the neuron *i*, we can rewrite (9) in terms of the objective function for the neuron model presented in (8), as given below

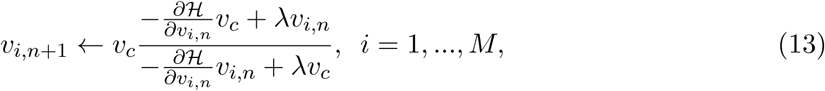

Where

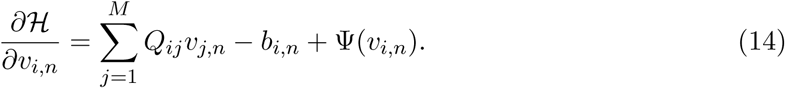

Then asymptotically from (12), and as shown in Appendix B, we have

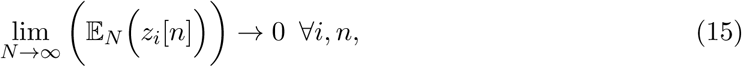

where 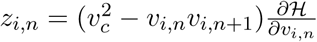. We first show the dynamics resulting from (13) for a trivial barrier function Ψ(.) = 0. Since *ℋ*(.) is a smooth function in this case, the neural variables *v*_*i,n*_ converge to a local minimum, such that

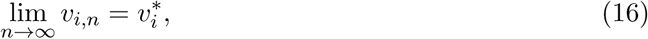

Therefore, (15) can be written as

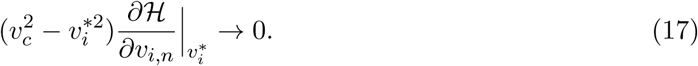

Thus as long as 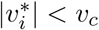, the gradient term goes to zero, ensuring that the dynamical system converges to the optimal solution within the domain defined by the bound constraints.

The dynamical system presented in (9) ensures that the steady-state neural responses 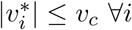. In the absence of the barrier term, the membrane potentials can converge to any value between −*v*_*c*_ and +*v*_*c*_ based on the effective inputs to individual neurons. Figure 2(a) illustrates this for 2 different neurons where **Q** is an identity matrix. For the sake of simplicity, we have considered the membrane potentials to be normalized in all the experiments in this paper (i.e., *v*_*c*_ = 1 *V*), and 0 *V* as the threshold voltage. Here 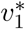 is hyperpolarized due to a negative stimulus, and 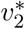 is depolarized beyond the threshold. Figure 2(b) shows the corresponding energy contours, where the steady-state neural responses encode the optimal solution of the energy function. We next show how this framework can be extended to a spiking neuron model when the trans-membrane current in the compartmental model described in (8) is approximated by a discontinuous Ψ(.). In general, the penalty function 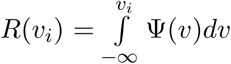 is chosen to be convex, where *R*(*v*_*i*_ *>* 0 *V*) *>* 0 *W* and *R*(*v*_*i*_ *≤* 0 *V*) = 0 *W*. Figure 2(c) shows one such form that has a gradient discontinuity at a threshold (*v*_*i*_ = 0 *V*) at which the neuron generates an action potential. The corresponding Ψ(.), also shown in Figure 2(c), is given by

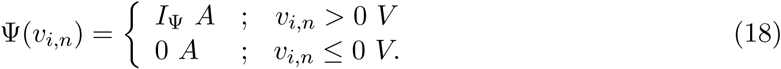

**Figure 2:**
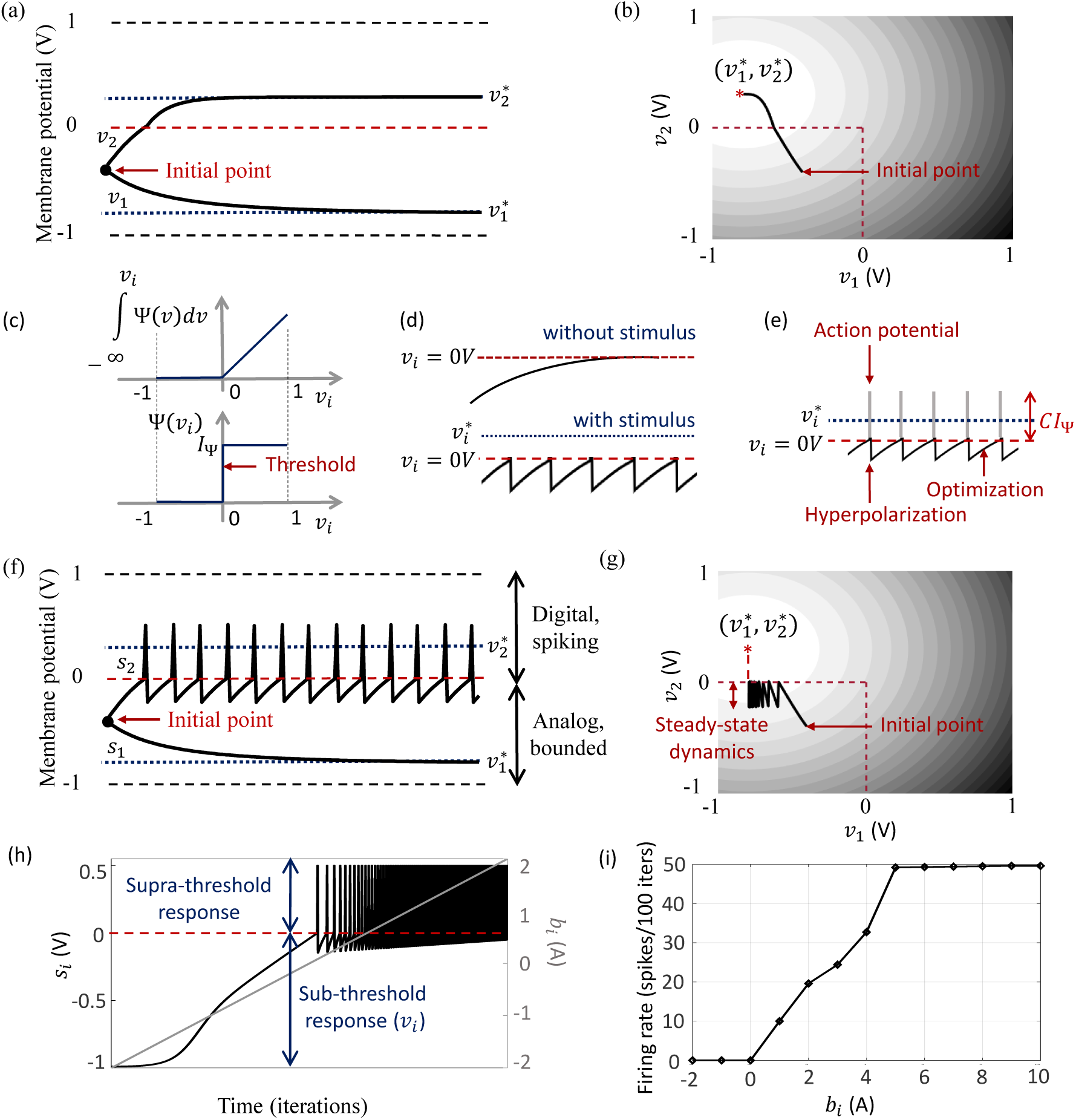
(a) Bounded dynamics in a 2-neuron network in absence of the barrier function; (b) Corresponding contour plot showing convergence of the membrane potentials in the presence of external stimulus; (c) The function ∫ Ψ(.)*dv* and its derivative Ψ(.) used in this paper for the spiking neuron model; (d) Time-evolution of the response *v*_*i*_ of a single neuron in the spiking model in the absence and presence of external stimulus; (e) The composite signal upon addition of spikes when *v*_*i*_ crosses the threshold; (f) Bounded and spiking dynamics in the same 2-neuron network in presence of barrier function; (g) Corresponding contour plot showing steady-state dynamics of the membrane potentials in the presence of external stimulus; (h) Plot of composite spike signal *s*_*i*_ of the spiking neuron model when the external current stimulus is increased; (i) Input-output characteristics for the spiking neuron model.

When there is no external stimulus *b*_*i*_, the neuron response converges to 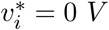 as in the non-spiking case, as illustrated in Figure 2(d) for a single neuron without any synaptic connections. When a positive stimulus *b*_*i*_ is applied, the optimal solution for *v*_*i*_, indicated by 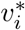, shifts upward to a level that is a function of the stimulus magnitude, also shown in Figure 2(d). However, a penalty term *R*(*v*_*i*_) of the form as described above works as a barrier function, penalizing the energy functional whenever *v*_*i*_ exceeds the threshold, thereby forcing *v*_*i*_ to reset below the threshold. The stimulus and the barrier function therefore introduce opposing tendencies, making *v*_*i*_ oscillate back and forth around the discontinuity (which, in our case, coincides with the threshold) as long as the stimulus is present. Thus when Ψ(.) is introduced, the potential *v*_*i,n*_ switches when Ψ(*v*_*i,n*_) *>* 0 *A* or only when *v*_*i,n*_ *>* 0 *V*. However, the dynamics of *v*_*i,n*_ remains unaffected for *v*_*i,n*_ *<* 0 *V*. During the brief period when *v*_*i,n*_ *>* 0 *V*, we assume that the neuron enters into a runaway state leading to a voltage spike. The composite spike signal *s*_*i,n*_, shown in Figure 2(e), is then treated as a combination of the sub-threshold and supra-threshold responses and is given by

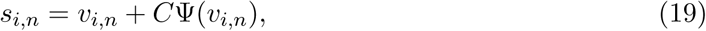

where the trans-impedance parameter *C >* 0 Ω determines the magnitude of the spike. An expectation over the composite signal *s*_*i,n*_ asymptotically encodes the new optimal solution 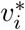 (dotted line), as shown in Figure 2(e) and later in Section 2.2.2. Note that in the current version, the proposed model does not explicitly model the runaway process that leads to the spike, unlike other neuron models [2, 3, 4]. However, it does incorporate the hyperpolarization part of the spike as a result of *v*_*i*_ oscillating around the gradient discontinuity. Thus a refractory period is automatically incorporated in between two spikes.

In order to show the effect of Ψ(.) on the nature of the solution, we plot the neural responses and contour plots for the 2-neuron network in Figures 2(f) and (g) for the same set of inputs as in Figures 2(a) and (b), when the barrier function is present. The penalty function produces a barrier at the thresholds, which are indicated by red dashed lines, transforming the time-evolution of *s*_2_ into a digital, spiking mode, where the firing rate is determined by the extent to which the neuron is depolarized. It can be seen from the neural trajectories in Figure 2(g) and from (8) that Ψ(.) *>* 0 behaves as a Lagrange parameter corresponding to the spiking threshold constraint *v*_*i,n*_ *<* 0.

In Appendix C, we outline how, for non-pathological cases, it can be shown from (12) that for spiking neurons, or for neurons whose membrane potentials *v*_*i,n*_ *>* −*v*_*c*_ ∀*n*,

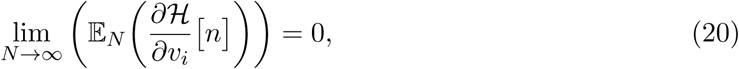

This implies that asymptotically the network exhibits limit-cycles about a single attractor or a fixed-point such that the time-expectations of its state variables encode this optimal solution. A similar stochastic first-order framework was used in [21] to derive a dynamical system corresponding to ΣΔ modulation for tracking low-dimensional manifolds embedded in high-dimensional analog signal spaces. Combining (14) and (20), we have

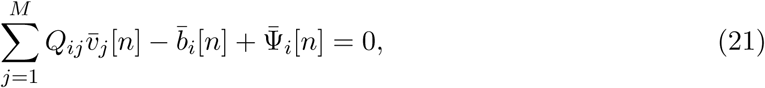

where 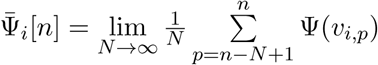. Rearranging the terms in (21), we obtain (7).

The penalty function 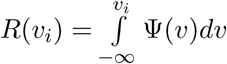 in the network energy functional in effect models the power dissipation due to spiking activity. For the form of *R*(.) chosen in this paper, the power dissipation due to spiking is taken to be zero below the threshold, and increases linearly above threshold. A plot of the composite spike signal for a ramp input for the spiking neuron model is presented in Figure 2(h). As *v*_*i,n*_ exceeds the threshold for a positive stimulus, the neuron enters a spiking regime and the firing rate increases with the input, whereas the sub-threshold response is similar to the non-spiking case. Figure 2(i) shows the tuning curve for the neuron as the input stimulus is increased. It is nearly sigmoidal in shape and shows how the firing rate reaches a saturation level for relatively high inputs. The proposed spiking neuron model based on the discrete-time Growth Transform dynamical system is summarized in Table 2.

**Table 2:**
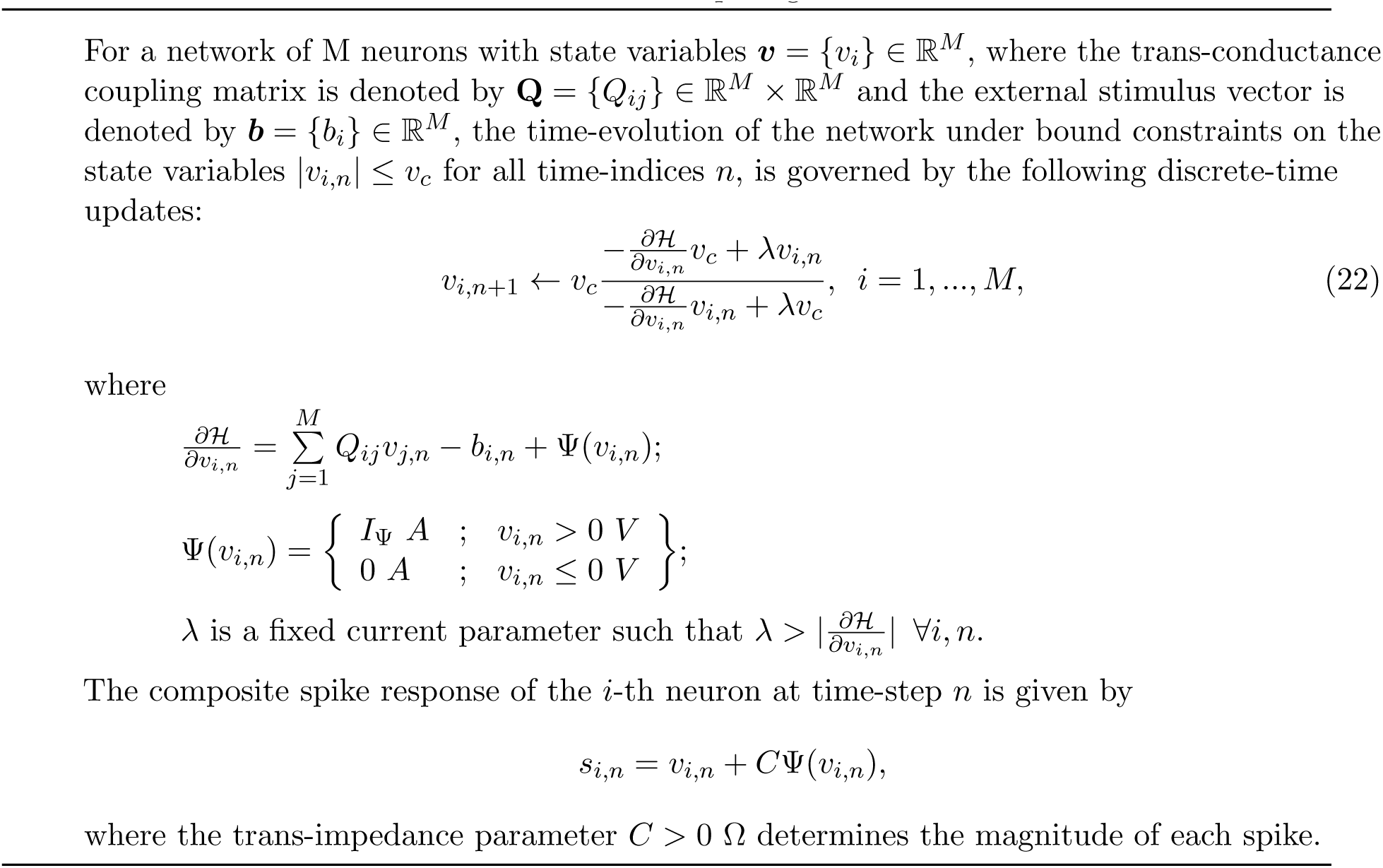
Discrete-time GT spiking neuron model

#### 2.2.2 Encoding stimuli as a combination of sub-threshold and supra-threshold dynamics

As explained previously, the penalty term *R*(*v*_*i*_) of the form presented above works analogous to a barrier function, penalizing the energy functional whenever *v*_*i,n*_ exceeds the threshold. This transforms the time-evolution of *v*_*i,n*_ into a spiking mode above the threshold, while keeping the sub-threshold dynamics similar to the non-spiking case. The Growth Transform dynamical system ensures that the membrane potentials are bounded, thereby implementing a squashing (compressive) function on the neural responses so that the network responses are bounded. We now show how the proposed model encodes external stimulus as a combination of spiking and bounded dynamics. In the steady-state, from (21) we can write

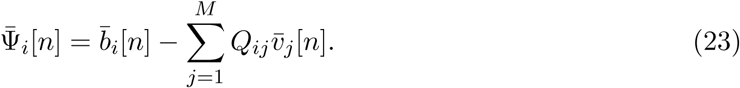

Thus the average spiking activity of the *i*-th neuron encodes the error between the average input and the weighted sum of membrane potentials. For a single, uncoupled neuron where

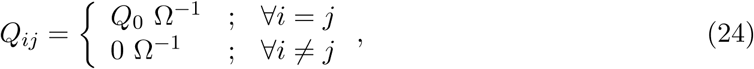

we have

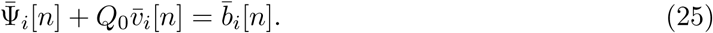

Multiplying (25) on both sides by *C* Ω, where we have chosen 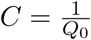, we have

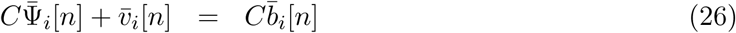

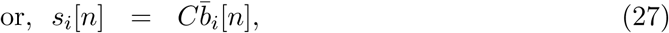

where we have used the relation (19). (27) indicates that through a suitable choice of the trans-impedance parameter *C*, the sum of sub-threshold and supra-threshold responses encodes the external input to the neuron. This is also the rationale behind adding a spike to the sub-threshold response *v*_*i,n*_, as illustrated in Figure 2(e), to yield the composite neural response. If *Q*_0_ = 0 Ω^−1^, we similarly have

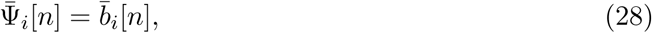

where the average spiking activity tracks the stimulus. Thus by defining the coupling matrix in various ways, we can obtain different encoding schemes for the network.

### 2.3 From neuron to network: Geometric interpretation of network dynamics

The remapping from standard coupled conditions of a spiking neural network to the proposed formulation admits a geometric interpretation of neural dynamics. Similar to the network coding framework presented in [8], we show in this section how the activity of individual neurons in a network can be visualized with respect to a network hyper-plane. This geometric interpretation can then be used to understand network dynamics in response to different stimuli.

Like a Hessian, if we assume that the matrix **Q** is positive-definite about a local attractor, there exists a set of vectors ***x***_*i*_ ∈ ℝ^*D*^, *i* = 1, …, *M* such that each of the elements *Q*_*ij*_ can be written as an inner product between two vectors as *Q*_*ij*_ = ***x***_*i*_.***x***_*j*_, 1 *≤ i, j ≤ M*. This is similar to kernel methods that compute similarity functions between pairs of vectors in the context of support vector machines [22]. This associates the *i*-th neuron in the network with a vector ***x***_*i*_, mapping it onto an abstract metric space R^*D*^ and essentially providing an alternate geometric representation of the neural network. From (21), the spiking activity of the *i*-th neuron for the *n*-th time-window can then be represented as

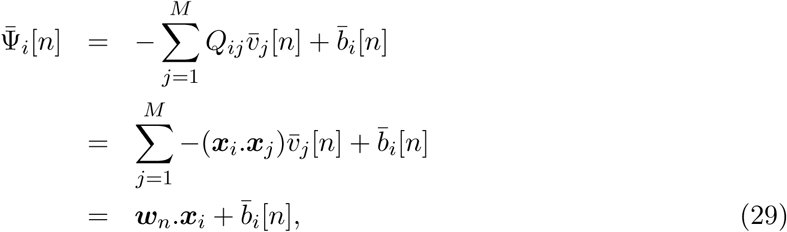

Where

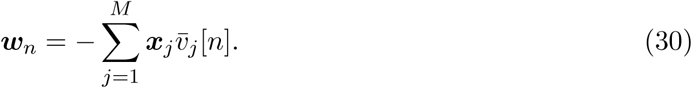

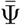 therefore represents the distance of the vector ***x***_*i*_ from a network hyperplane in the *D*-dimensional vector space, which is parameterized by the weight vector ***w***_*n*_ and offset 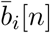. When a stimulus 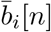 is applied, the hyperplane shifts, leading to a stimulus-specific value of this distance for each neuron that is also dependent on the network configuration **Q**. Hence, 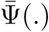 is denoted as a ‘network variable’, that signifies how the response of each neuron is connected to the rest of the network. Note that we can also write the synaptic weight elements in a kernelized form as *Q*_*ij*_ = *K*(***x***_*i*_).*K*(***x***_*j*_), where *K*(.) is a nonlinear transformation function, defining a non-linear boundary for each neuron. This idea of a dynamic and stimulus-specific hyperplane can offer intuitive interpretations about several population dynamics reported in literature and have been elaborated on in the Results section.

### 2.4 Complete continuous-time model of the Growth Transform neuron

Single neurons show a vast repertoire of response characteristics and dynamical properties that lend richness to their computational properties at the network level. Izhikevich in [23] provides an extensive review of different spiking neuron models and their ability to produce the different dynamics observed in biology. In this section, we extend the proposed model into a continuous-time dynamical system, which enables us to reproduce a vast majority of such dynamics and also allows us to explore interesting properties in coupled networks. In Appendix D, we derive the continuous-time version of the dynamical system using a special property of Growth Transforms. The complete neuron model is summarized in Table 3.

**Table 3:**
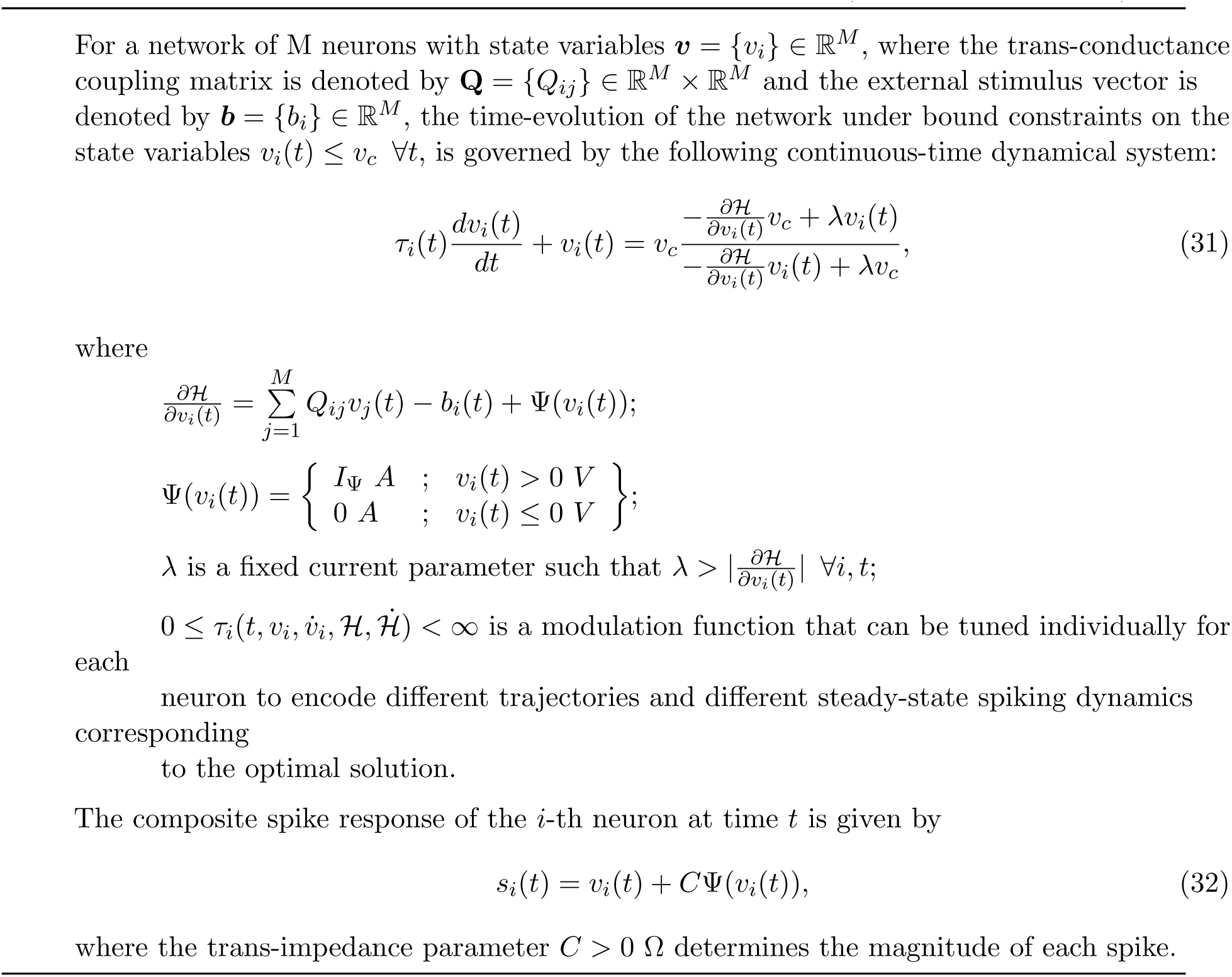
Complete continuous-time GT spiking neural network (Proof in Appendix D)

The operation of the proposed neuron model is therefore governed by two sets of dynamics: (a) minimization of the network energy functional *ℋ*; (b) modulation of the trajectory using a time-constant *τ*_*i*_(*t*), also referred to as modulation function in this paper. Fortunately, the evolution of *τ*_*i*_(*t*) can be made as complex as possible without affecting the asymptotic fixed-point solution of the optimization process. It can be a made a function of local variables like *v*_*i*_ and 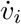 or a function of global/network variables like *H* and 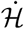. Different choices of the modulation function can lead to different trajectories followed by the neural variables under the same energy contour, as illustrated in Figure 3. In the Results section, we show how different forms of *τ*_*i*_(*t*) produce different sets of neuronal dynamics consistent with the dynamics that have been

**Figure 3:**
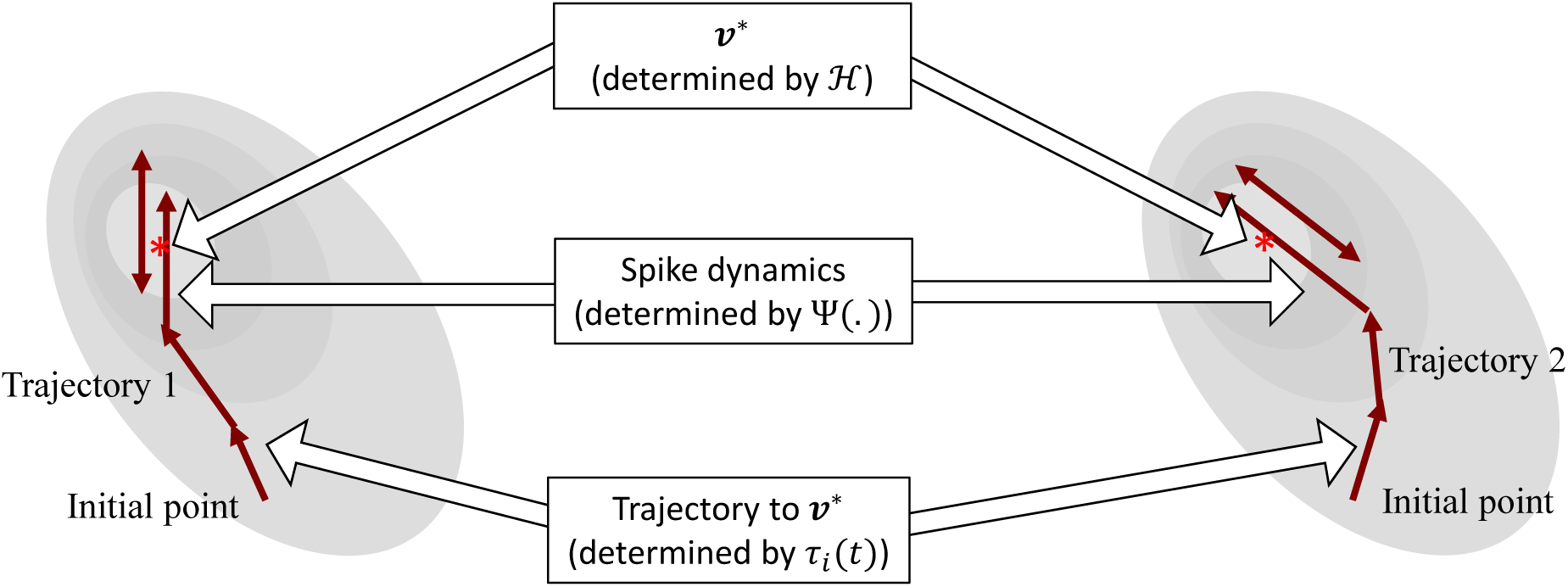
Decoupling of network solution, spike shape and response trajectory using the proposed model. Different modulation functions lead to different steady-state spiking dynamics under the same energy contour.

## 3 Results

The proposed approach enables us to decouple the three following aspects of the spiking neural network:

a. Fixed points of the network energy functional, which depend on the network configuration and external inputs;
b. Nature and shape of neural responses, without affecting the network minimum; and
c. Spiking statistics and transient neural dynamics at the cellular level, without affecting the network minimum or spike shapes.

This makes it possible to independently control and optimize each of these neuro-dynamical properties without affecting the others. The first two aspects arise directly from an appropriate selection of the energy functional and were demonstrated in Section 2.2.1. In this section, we show how the modulation function, in essence, loosely models cell excitability, and can be varied to tune transient firing statistics based on local and/or global variables. This allows us to encode the same optimal solution using widely different firing patterns across the network, and have unique potential benefits for neuromorphic applications. Codes for the representative examples given in this section are available at [11].

### 3.1 Single-neuron dynamics

We first show how we can reproduce a number of single-neuron response characteristics by changing the modulation function *τ*_*i*_(*t*) in the neuron model. For this, we consider an uncoupled network, where

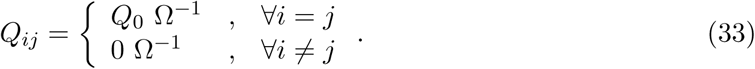

We will subsequently extend these dynamics to build coupled networks with interesting properties like memory and global adaptation for energy-efficient neural representation. The results reported here are representative of the types of dynamical properties the proposed model can exhibit, but are by no means exhaustive. Readers are encouraged to experiment with different inputs and network parameters in the software (MATLAB^©^) implementation of the Growth Transform neuron model [11]. The tool enables users to visualize the effects of different modulation functions and other parameters on the neural dynamics, as well as the time-evolution of population trajectories and the network energy function with different inputs and under different initial conditions.

#### 3.1.1 Standard tonic-spiking response

When stimulated with a constant current stimulus *b*_*i*_, a vast majority of neurons fire single, repetitive action potentials for the duration of the stimulus, with or without adaptation [24, 25, 26]. The proposed model shows tonic spiking without adaptation when the modulation function *τ*_*i*_(*t*) = *τ*, where *τ >* 0 s. A simulation of tonic spiking response using the neuron model is given in Figure 4(a).

**Figure 4:**
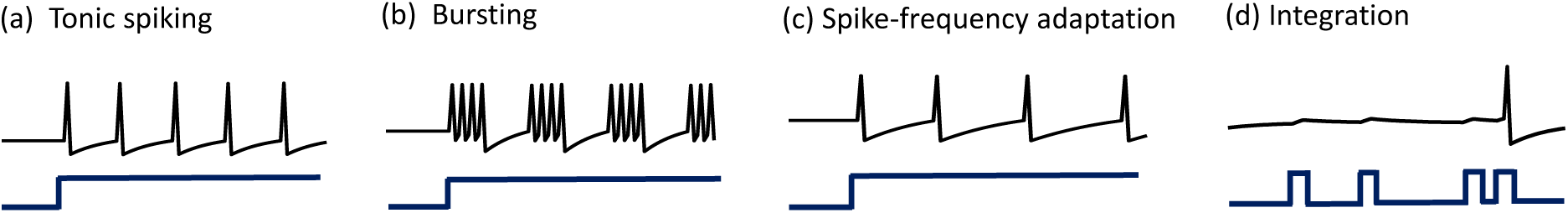
(a)-(d) Simulations demonstrating different single-neuron responses obtained using the GT neuron model.

#### 3.1.2 Bursting response

Bursting neurons fire discrete groups of spikes interspersed with periods of silence in response to a constant stimulus [24, 25, 27, 28]. Bursting arises from an interplay of fast ionic currents responsible for spiking, and slower intrinsic membrane currents that modulate the spiking activity, causing the neuron to alternate between activity and quiescence. Bursting response can be simulated in the proposed model by modulating *τ*_*i*_(*t*) at a slower rate compared to the generation of action potentials, in the following way

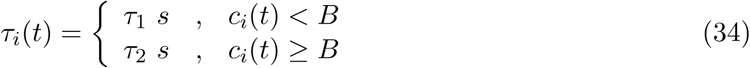

where *τ*_1_ *> τ*_2_ *>* 0 s, *B* is a parameter and the count variable *c*_*i*_(*t*) is updated according to

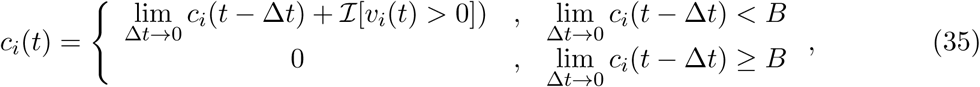

*ℑ*[.] being an indicator function. Simulation of a bursting neuron in response to a step input is given in Figure 4(b).

#### 3.1.3 Spike-frequency adaptation

When presented with a prolonged stimulus of constant amplitude, many cortical cells initially respond with a high-frequency spiking that decays to a lower steady-state frequency [29]. This adaptation in the firing rate is caused by a negative feedback to the cell excitability due to the gradual inactivation of depolarizing currents or activation of slow hyperpolarizing currents upon depolarization, and occur at a time-scale slower than the rate of action potential generation. We modeled spike-frequency adaptation by varying the modulation function according to

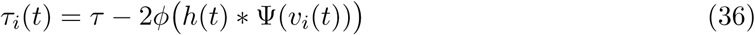

where *h*(*t*) * Ψ(*v*_*i*_)(*t*) is a convolution operation between a continuous-time first-order smoothing filter *h*(*t*) and the spiking function Ψ(*v*_*i*_(*t*)), and

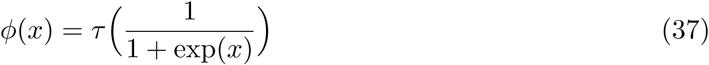

is a compressive function that ensures 0 *≤ τ*_*i*_(*t*) *≤ τ* s. The parameter *τ* determines the steady-state firing rate for a particular stimulus. A tonic-spiking response with spike-frequency adaptation is shown in Figure 4(c).

#### 3.1.4 Integrator response

When the baseline input is set slightly negative so that the fixed point is below the threshold, the neuron works like a leaky integrator as shown in Figure 4(d), preferentially spiking to high-frequency or closely-spaced input pulses that are more likely to make *v*_*i*_ cross the threshold.

### 3.2 Coupled spiking network with pre-synaptic adaptation

We can extend the proposed framework to a network model where the neurons, apart from external stimuli, receive inputs from other neurons in the network. We begin by considering **Q** to be a positive-definite matrix, which gives a unique solution of (8). Although elements of the coupling matrix **Q** already capture the interactions among neurons in a coupled network, we can further define the modulation function as follows to make the proposed model behave as a standard spiking network

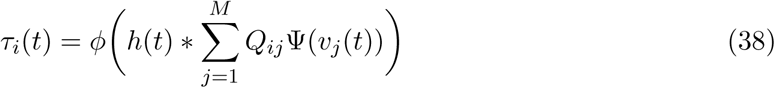

with the compressive-function *φ*(.) given by (37). (38) ensures that *Q*_*ij*_ *>* 0 corresponds to an excitatory coupling from the pre-synaptic neuron *j*, and *Q*_*ij*_ *<* 0 corresponds to an inhibitory coupling, as demonstrated in Figure 5(a). Note that irrespective of whether such a pre-synaptic adaptation is implemented or not, the neurons under the same energy landscape would converge to the same sub-domain, albeit with different response trajectories and steady-state limit-cycles. This is illustrated in Figure 5(b) which plots the energy contours for a two-neuron network corresponding to a **Q** matrix with excitatory and inhibitory connections and a fixed stimulus vector ***b***. Figure 5(b) also shows the responses of the two neurons starting from the same initial conditions, with and without pre-synaptic adaptation (where the latter corresponds to the case where the only coupling between the two neurons is through the coupling matrix **Q**, but there is no pre-synaptic spike-time dependent adaptation). Because the energy landscape is the same in both cases, the neurons converge to the same sub-domain, but with widely varying trajectories and steady-state response patterns.

**Figure 5:**
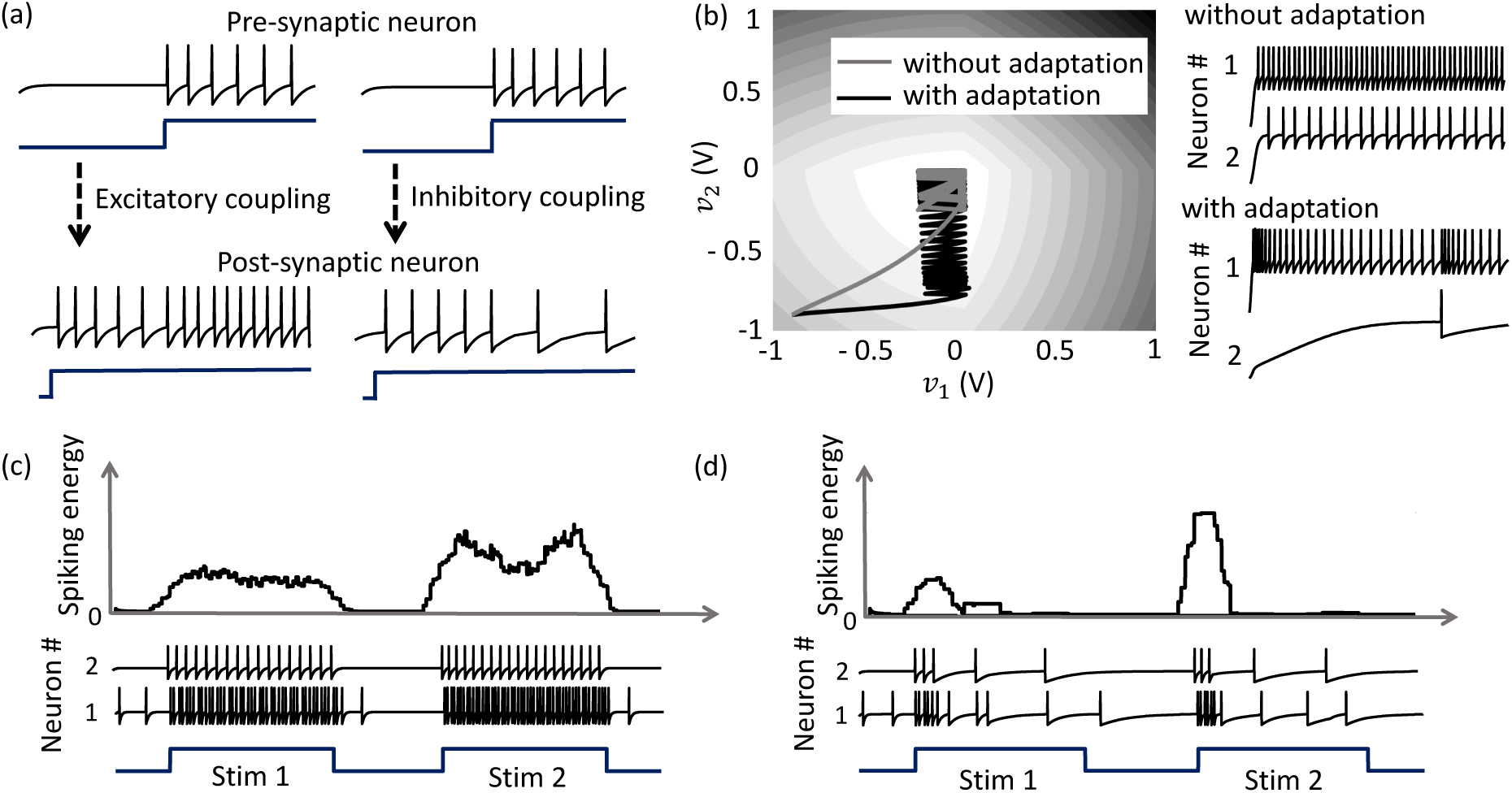
(a) Results from a 2-neuron network with excitatory and inhibitory couplings; (b) Energy optimization process under different conditions lead to different limit cycles within the same energy landscape. (c)-(d) Mean spiking energy ∫ Ψ(.)*dv* and firing patterns in response to a step input without and with global adaptation respectively.

### 3.3 Coupled network with pre-synaptic and global adaptation

Apart from the pre-synaptic adaptation that changes individual firing rates based on the input spikes received by each neuron, neurons in the coupled network can be made to adapt according to the global dynamics by changing the modulation function as follows

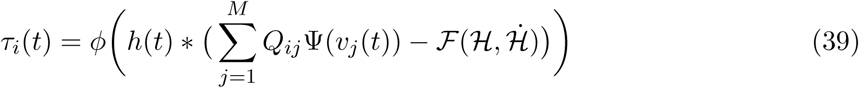

with the compressive-function *φ*(.) given by (37). The new function *ℱ* (.) is used to capture the dynamics of the network cost-function. As the network starts to stabilize and converge to a fixed-point, the function *τ*_*i*_(.) adapts to reduce the spiking rate of the neuron without affecting the steady-state solution. Figures 5(c) and (d) shows the time-evolution of the spiking energy ∫ Ψ(.)*dv* and the spike-trains for a two-neuron network without global adaptation and with global adaptation respectively, using the following form for the adaptation term

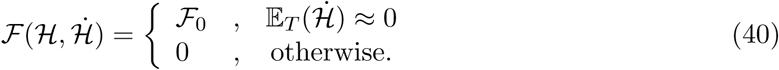

where *ℱ*_0_ *>* 0 is a tunable parameter. This feature is important in designing energy-efficient spiking networks where energy is only dissipated during transients.

### 3.4 Network response and network trajectories

In order to outline the premises of the next few experiments on population dynamics using the geometric interpretation outlined in Section 2.3, we consider a small network of neurons on a two-dimensional co-ordinate space, and assign arbitrary inputs to the neurons. A Gaussian kernel is chosen for the coupling matrix **Q** as follows

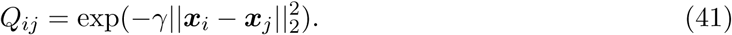

This essentially clusters neurons with stronger couplings between them closer to each other on the co-ordinate space, while placing neurons with weaker couplings far away from each other. A network consisting of 20 neurons is shown in Figure 6(a), which also shows how the spiking activity changes as a function of the location for the particular network configuration and input stimulus vector. Each neuron is color coded based on the mean firing rate (normalized w.r.t. the maximum mean firing rate) with which it responds when the stimulus is on. Figure 6(b) shows the spike raster for the entire network. We see that the responsiveness of the neurons to a particular stimulus increases with the distance at which it is located from the hypothetical hyperplane in the high-dimensional space to which the neurons are mapped through kernel transformation. We show below how this geometric representation can provide insights on population-level dynamics in the network considered.

**Figure 6:**
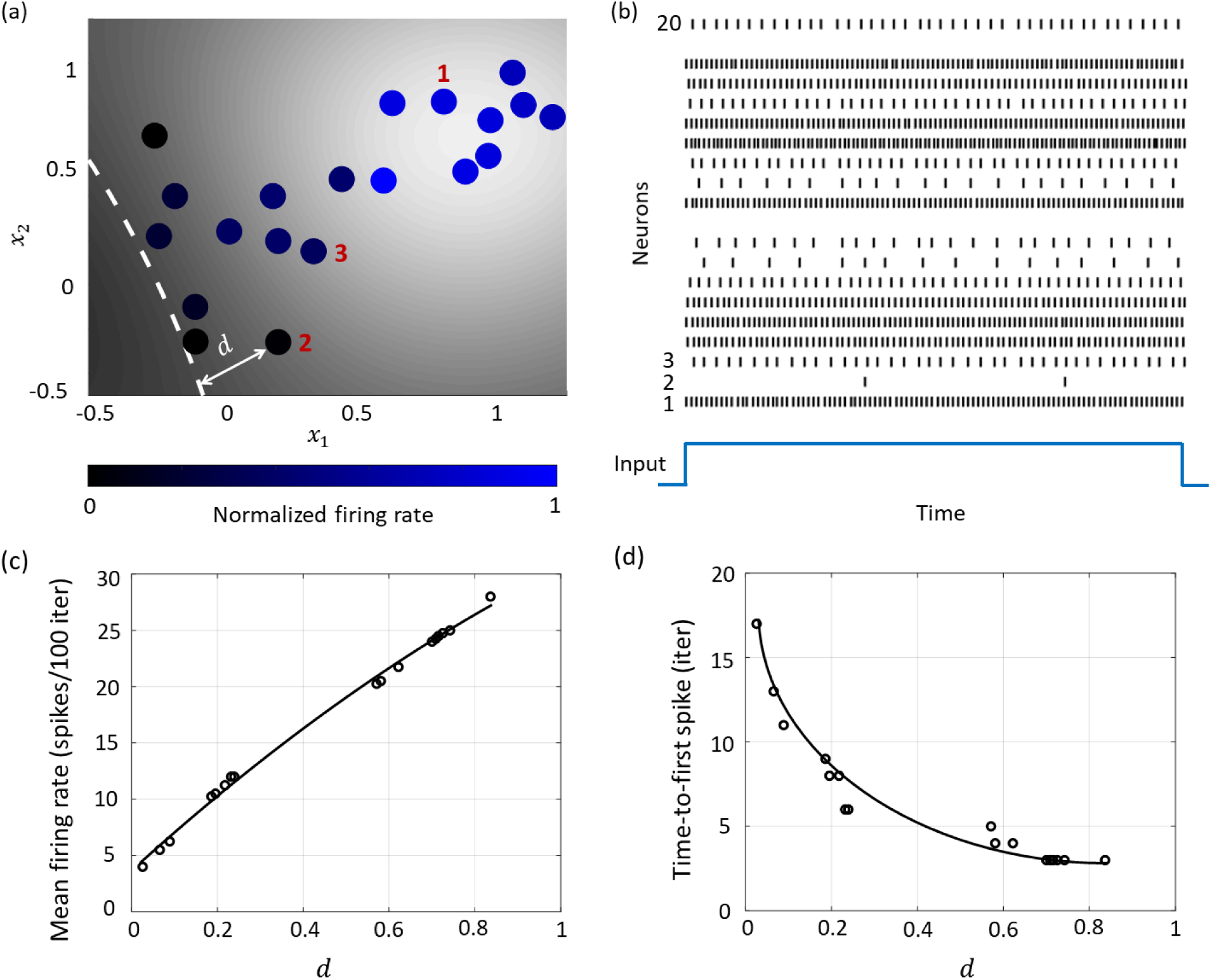
(a) Contour plot of spiking activity corresponding to a particular stimulus vector. Neurons are colored according to their mean firing rate (normalized w.r.t. the maximum firing rate) during the stimulus period. The white dashed line is the contour corresponding to 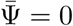. (b) Spike raster for all neurons for the input in (a). (c) The mean firing rate and (d) time-to-first spike as a function of the distance *d* for each neuron in the network.

#### 3.4.1 Rate and temporal coding

The Growth Transform neural network inherently shows a number of encoding properties that are commonly observed in biological neural networks [13, 30]. For example, the firing rate averaged over a time window is a popular rate coding technique that claims that the spiking frequency or rate increases with stimulus intensity [31]. A temporal code like the time-to-first-spike posits that a stronger stimulus brings a neuron to the spiking threshold faster, generating a spike, and hence relative spike arrival times contain critical information about the stimulus [32].

These coding schemes can be interpreted under the umbrella of network coding using the same geometric representation as considered above. Here, the responsiveness of a neuron is closely related to its proximity to the hyperplane. The neurons which exhibit more spiking are located at a greater distance from the hyperplane. We see from Figures 6(c) and (d) that as this value increases, the average firing rate of a neuron (number of spikes in a fixed number of time-steps or iterations) increases, and the time-to-first spike becomes progressively smaller. Neurons with a distance value below a certain threshold do not spike at all during the stimulus period, and therefore have a mean firing rate of zero and time-to-spike at infinity. Therefore, based on how the network is configured in terms of synaptic inputs and connection strengths, the spiking pattern of individual neurons conveys critical information about the boundary and their placement with respect to it.

#### 3.4.2 Network coding and neural population trajectories

The encoding of a stimulus in the spatiotemporal evolution of activity in a large population of neurons is often represented in neurobiological literature by a unique trajectory in a high-dimensional space, where each dimension accounts for the time-binned spiking activity of a single neuron. Projection of the high-dimensional activity to two or three critical dimensions using dimensionality reduction techniques like Principal Component Analysis (PCA) and Linear Discriminant Analysis (LDA) have been widely used across organisms and brain regions to shed light on how neural population response evolves when a stimulus is delivered [33, 34]. For example in identity coding, trajectories corresponding to different stimuli evolve towards different regions in the reduced neural subspace, that often become more discriminable with time and are stable over repeated presentations of a particular stimulus [33, 34, 35]. We show how this can be explained in the context of the geometric interpretation.

For the same network as above, we start with the simplest possible experiment, starting from the same baseline, and perturbing the stimulus vector in two different directions. This pushes the boundary in two different directions, exciting different subsets of neurons, as illustrated in Figures 7(a) and (b). A similar dimensionality reduction to three principal components in Figure 7(c) shows the neural activity unfolding in distinct stimulus-specific areas of the neural subspace. The two contour plots also show that some neurons may spike for both the inputs, while some spike selectively for one of them. Yet others may not show any spiking for either stimulus, but may spike for some other stimulus vector and the corresponding stimulus-specific boundary.

**Figure 7:**
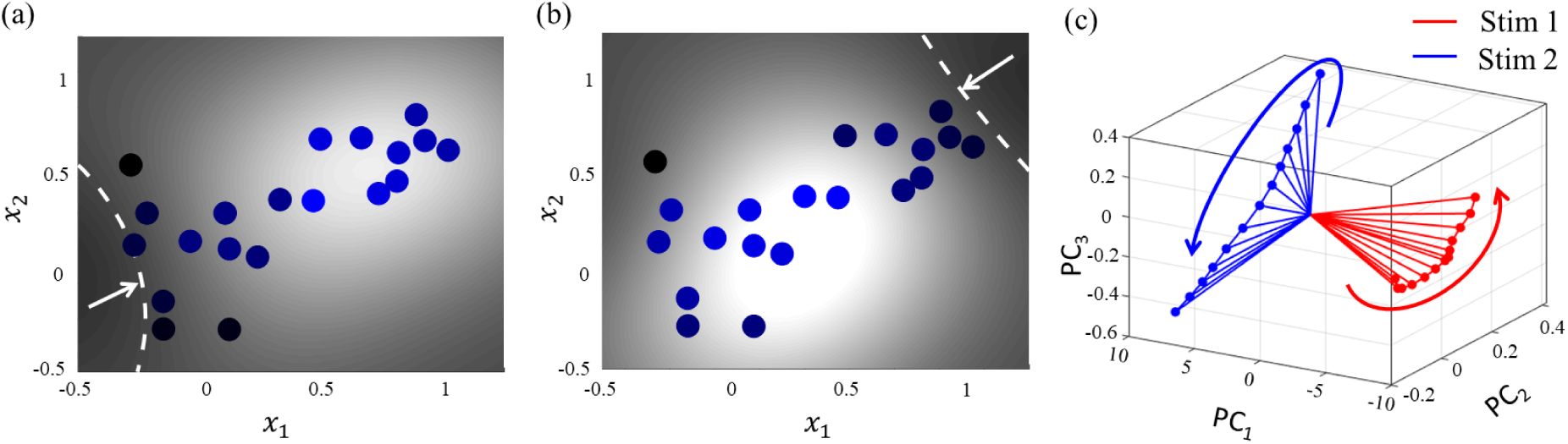
(a) and (b) Perturbation of the stimulus vector in different directions for the same network produces two different contours. (c) Corresponding population activities trace different trajectories in the neural subspace.

### 3.5 Coupled spiking network with non-positive definite Q

As illustrated in Figure 8, a coupled spiking network can function as a memory element, when **Q** is a non-positive definite matrix and

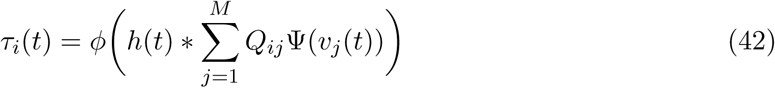

due to the presence of more than one attractor state. We demonstrate this by considering two different stimulus histories in a network of four neurons, where a stimulus ‘Stim 1a’ precedes another stimulus ‘Stim 2’ in Figures 8(a), (c) and (e), and a different stimulus ‘Stim 1b’ precedes ‘Stim 2’ in Figures 8(b), (d) and (f). Here, each ‘stimulus’ essentially corresponds to a different input vector ***b***. For an uncoupled network, where neurons do not receive any inputs from other neurons, the network energy increases when the first stimulus is applied and returns to zero afterwards, and the network begins from the same state again for the second stimulus as for the first, leading to the same firing pattern for the second stimulus, as shown in Figures 8(a) and (b), independent of the history. For a coupled network with a positive definite coupling matrix **Q**, reinforcing loops of spiking activity in the network may not allow the network energy to go to zero after the first stimulus is removed, and the residual energy may cause the network to exhibit a baseline activity that depends on stimulus history, as long as there is no dissipation. When the second stimulus is applied, the initial conditions for the network are different for the two stimulus histories, leading to two different transients until the network settles down into the same steady-state firing patterns, as shown in Figures 8(c) and (d). For a non-positive definite coupling matrix **Q** however, depending on the initial condition, the network may settle down to different solutions for the same second stimulus, due to the possible presence of more than one local minimum. This leads to completely different transients as well as steady-state responses for the second stimulus, as shown in Figure 8(e) and (f). This history-dependent stimulus response could serve as a short-term memory, where residual network energy from a previous external input subserves synaptic interactions among a population of neurons to set specific initial conditions for a future stimulus based on the stimulus history, forcing the network to settle down in a particular attractor state.

**Figure 8:**
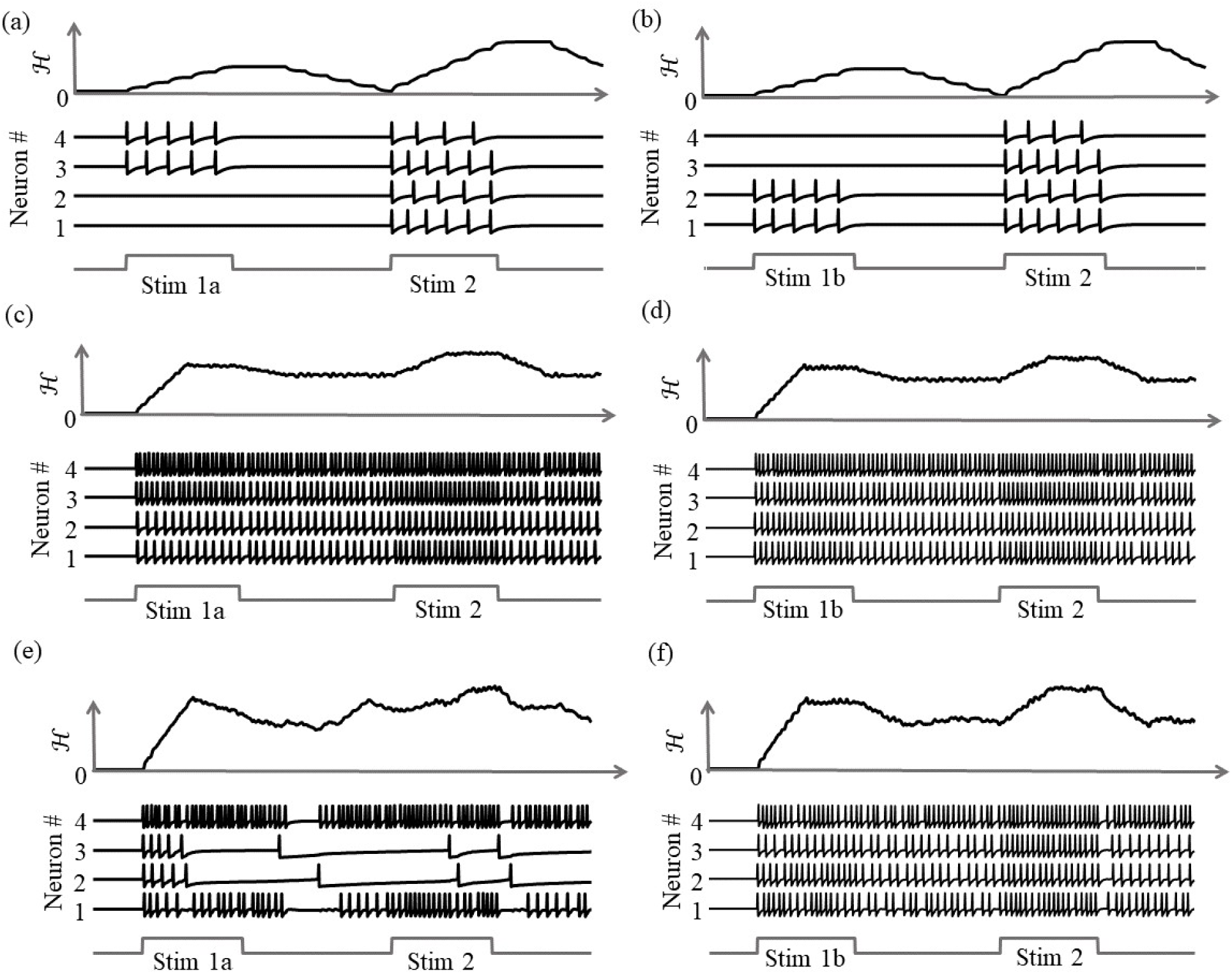
Stimulus response for a 4-neuron network with different stimulus histories for: (a)-(b) an uncoupled network; (c)-(d) a coupled network with a positive definite coupling matrix **Q**; (e)-(f) a coupled network with a non-positive definite coupling matrix **Q**.

### 3.6 Associative memory network using Growth Transform neuron models

Associative memories are neural networks which can store memory patterns in the activity of neurons in a network through a Hebbian modification of their synaptic weights; and recall a stored pattern when stimulated with a partial fragment or a noisy version of the pattern [36]. Various works have studied associative memories using networks of spiking neuron models having different degrees of abstraction and architectural complexities [37, 38]. Here, we demonstrate using an associative memory network of Growth Transform neurons how we can use network trajectories to recall stored patterns, and moreover, use global adaptation to do so using very few spikes and high recall accuracy.

Our network comprises *M* = 100 neurons, out of which a randomly selected subset *m* = 10 are active for any stored memory pattern. The elements of the transconductance coupling matrix are set according to the following standard Hebbian learning rule

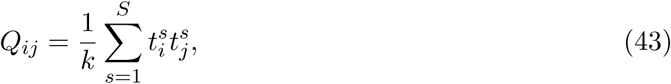

where *k* is a scaling factor and ***t***^*s*^ ∈ [0, 1]^*M*^, *s* = 1, …, *S*, are the binary patterns stored in the network. During the recall phase, only half of the cells active in the original memory are stimulated with a steady depolarizing input, and the spiking pattern across the network is recorded. Instead of determining the active neurons during recall through thresholding and directly comparing with the stored binary pattern, we quantitatively measure the recall performance of the network by computing the mean distance between each pair of original-recall spiking dynamics as they unfold over time. This ensures that we not only take into account the firing of the neurons that belong to the pattern albeit are not directly stimulated, but also enables us to exploit any contributions from the rest of the neurons in making the spiking dynamics more dissimilar in comparison to recalls for other patterns.

When the network is made to globally adapt according to the system dynamics, the steady-state trajectories can be encoded using very few spikes. Figures 9 (a) and (b) show the raster plots for the stored patterns without and with global adaptation respectively when *S* = 10; and Figures 9 (c) and (d) are the corresponding plots during recall. For each recall pattern, spike patterns for the directly stimulated neurons are plotted first, followed by the other 5 neurons that are not directly stimulated but belong to the pattern; and finally the rest of the neurons in random order. The ordering of neurons is kept the same for plotting spike rasters for the stored patterns. During decoding, a straightforward metric using the average distance between time-binned mean firing rates for the original and recall trajectories produces similarity matrices presented in Figures 10(a) and (b), where we see that global adaptation does not perform as well. However, the information in this case also lies in the spike-times and changes in firing rate over time for each neuron. Including these features in the decoding vectors for stored and recalled patterns, we get clean recalls in both cases as shown in Figures 10(c) and (d). The decoding vector for the *n*-th time-bin in this case is given by

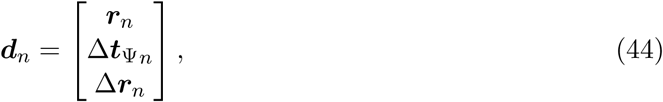

where ***r***_*n*_, Δ***t***_Ψ*n*_ and Δ***r***_*n*_ are the vectors of mean firing rates, mean inter-spike intervals and changes in the mean firing rates for the *n*-th bin for the entire network respectively. The mean inter-spike interval is set equal to the bin length if there is a single spike over the entire bin length, and equal to twice the bin length if there are none. Note that the inter-spike interval computed for one time-bin may be different from (1*/r*), particularly for low firing rates, and hence encodes useful information. The similarity metric between the *u*-th stored pattern and the *v*-th recall pattern is given by

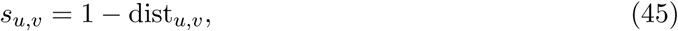

where dist_*u,v*_ is the mean Euclidean distance between the two decoding vectors over the total number of time-bins, normalized between [0, 1].

**Figure 9:**
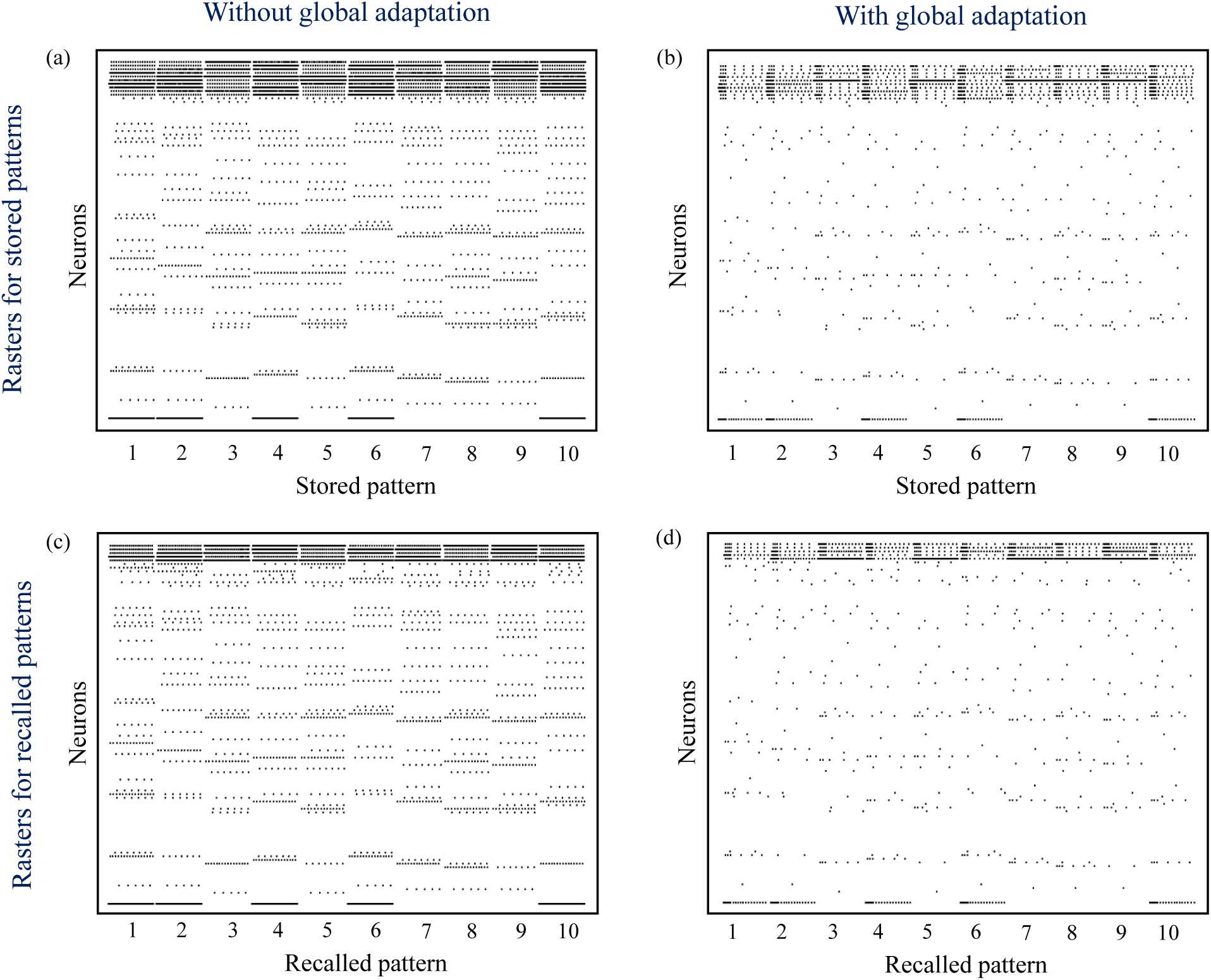
(a)-(b) Spike rasters for the 10 stored patterns without and with global adaptation respectively; (c)-(d) Spike rasters for the 10 recall cases without and with global adaptation respectively.

**Figure 10:**
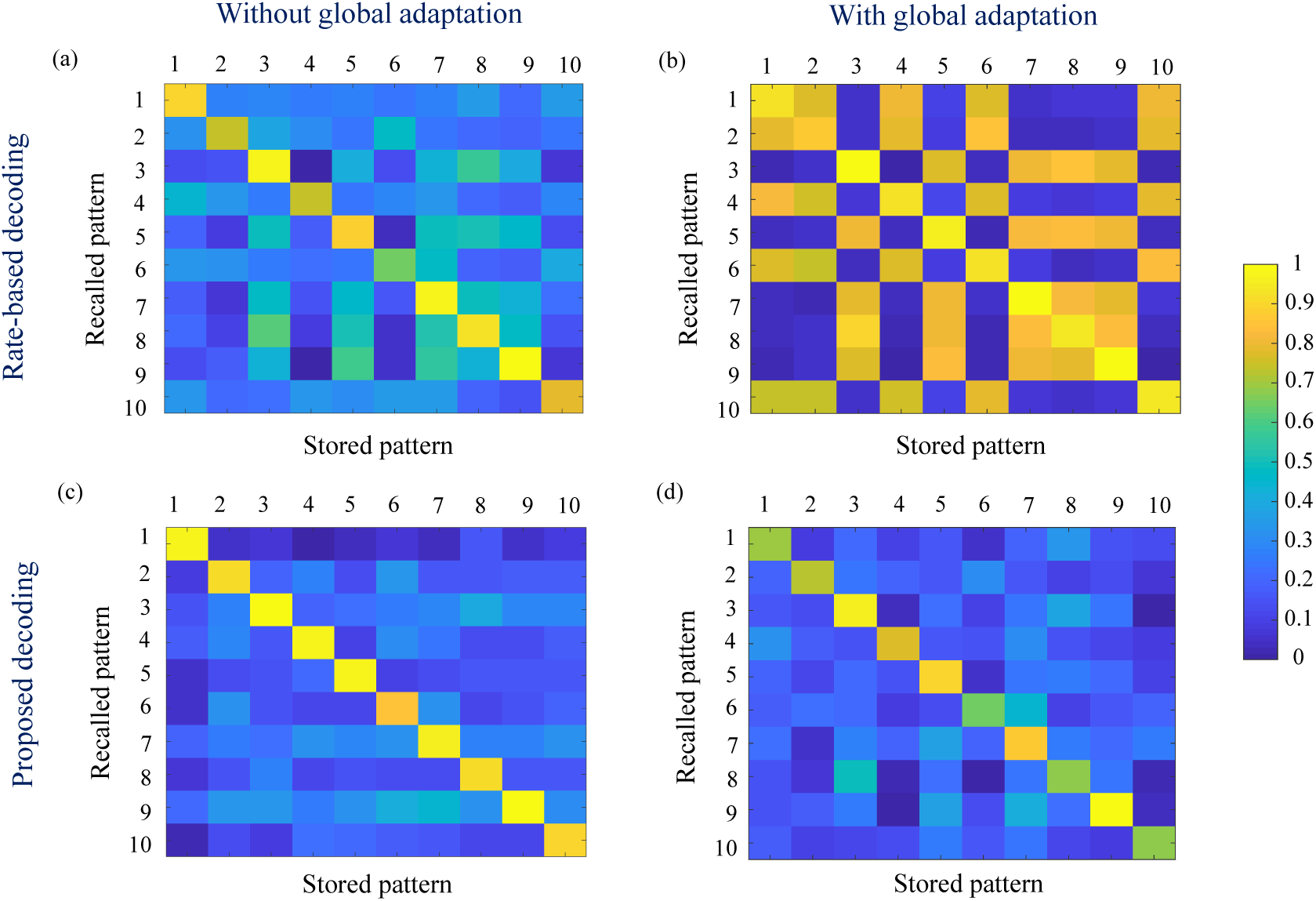
(a)-(b) Similarity matrices between storage and recall with a rate-based decoding metric; (c)-(d) Similarity matrices with a decoding metric that also includes spike-times and changes in mean firing rate.

To estimate the capacity of the network, we calculate the mean recall accuracy over 10 trials for varying number of stored patterns, both with and without global adaptation. Figure 11(a) plots the mean recall accuracy for different number of patterns stored for the two cases, and Figure 11(b) plots the mean number of spikes for each storage. For each plot, the shaded region indicates the range of values across trials. As expected, the accuracy is 100% for lesser storage,but degrades with higher loading. However with global adaptation, the degradation is seen to be more graceful for a large range of storage with the decoding used in Figures 10(c) and (d), allowing the network to recall patterns more accurately using much fewer spikes. Hence by exploiting suitable decoding techniques, we can implement highly energy-efficient spiking associative memory networks with high storage capacity.

**Figure 11:**
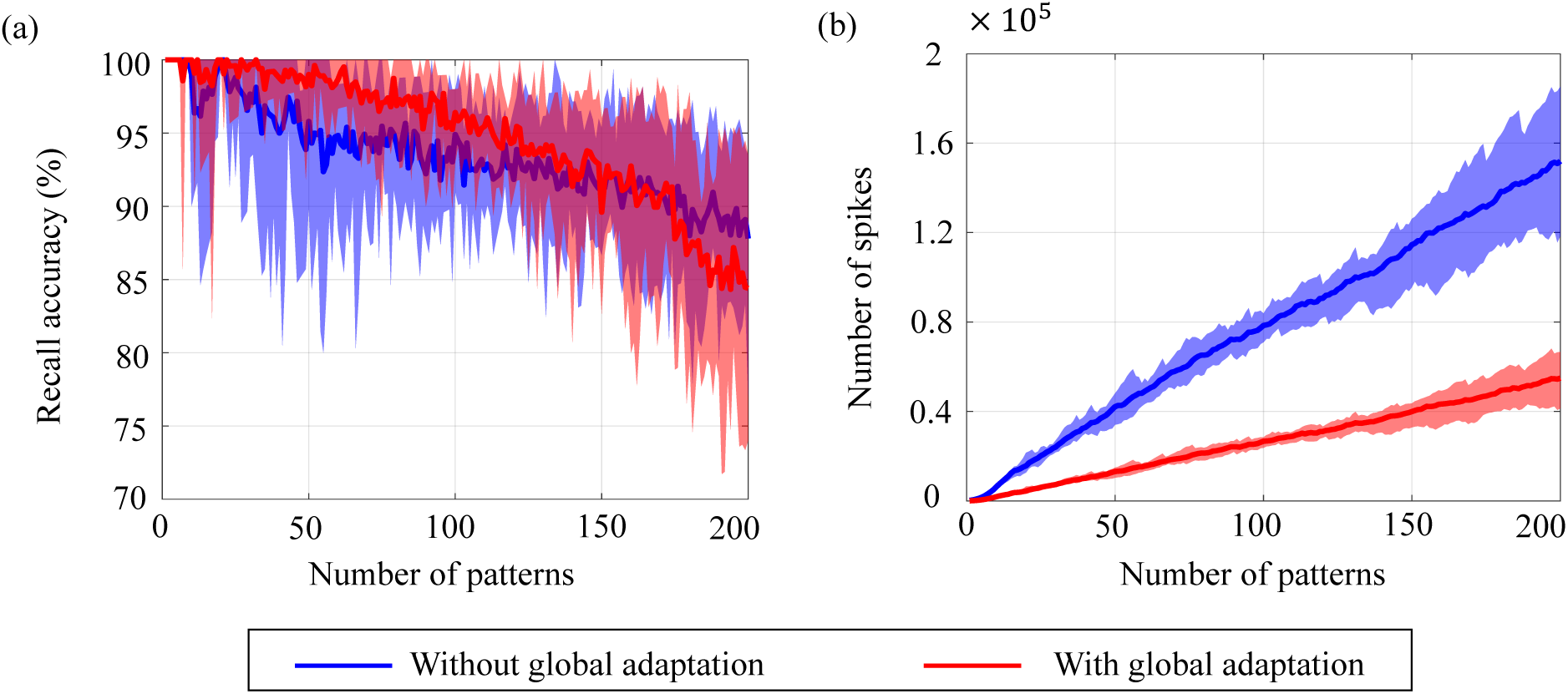
Ensemble plots showing (a) mean recall accuracy and (b) mean number of spikes as memory load increases for the network, in the absence as well as presence of global adaptation. The range of values across the ensemble is shown by the shaded area.

Note that the recall accuracy using global adaptation deteriorates faster for *>* 175 patterns. The proposed decoding algorithm, which determines the recall accuracy, takes into account the mean spiking rates, inter-spike intervals and changes in spike rates. It is possible that as the number of spikes is reduced through the use of global adaption, the information encoded in first-order differences (inter-spike intervals or spike rates) may not be sufficient to encode information at high fidelity, resulting in the degradation in recall accuracy when the number of patterns increased. However, augmenting the decoding features with higher-order differences in inter-spike intervals or spike rates may lead to an improved performance for higher storage.

#### 3.6.1 Classification of noisy MNIST images

Aside from pattern completion, associative networks are also commonly used for identifying patterns from their noisy counterparts. We use a similar associative memory network as above to classify images from the MNIST dataset which were corrupted with additive white Gaussian noise at different signal-to-noise ratios (SNRs), and which were, unlike in the previous case, unseen by the network before the recall phase. The network size in this case was *M* = 784, the number of pixels in each image, and the connectivity matrix was set using a separate, randomly selected subset of 5000 binary, thresholded images from the training dataset according to (43). Unseen images from the test dataset were corrupted at different SNRs and fed to the network after binary thresholding. Figures 12(a)-(c) shows an example of a test image at different SNRs after binary thresholding. As before, the non-zero pixels got a steady depolarizing input. A noisy test image was assigned to the class corresponding to the closest training image according to the similarity metric in (45).

**Figure 12:**
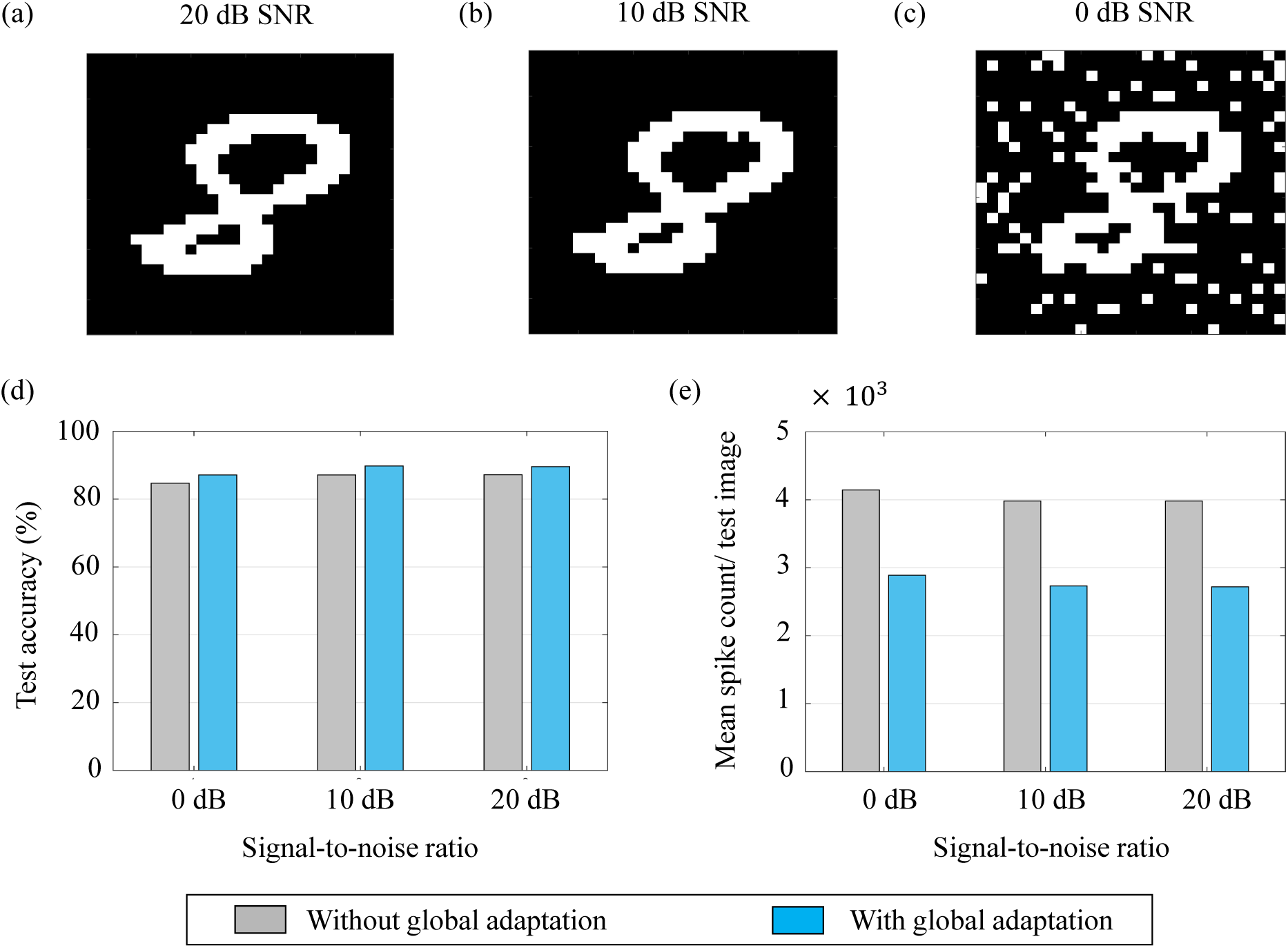
(a), (b) and (c) An example of a test image corrupted with additive white Gaussian noise at 20 dB, 10 dB and 0 dB SNR respectively; (d) and (e) Test accuracy and mean spike count/test image for different noise levels.

The test accuracies and mean spike counts for a test image are plotted in Figures 12(d) and (e) respectively for different noise levels. We see that even for relatively high noise levels, the network has a robust classification performance. As before, a global adaptation based on the state of convergence of the network produces a slightly better performance with fewer spikes per test image.

## 4 Conclusions

This paper introduces the theory behind a new spiking neuron and population model based on the Growth Transform dynamical system. The system minimizes an appropriate energy functional under realistic physical constraints to produce emergent spiking activity in a population of neurons. The proposed work is the first of its kind to treat the spike generation and transmission processes in a spiking network as an energy-minimization problem involving continuous-valued neural state variables like the membrane potential. The neuron model and its response are tightly coupled to the network objective, and is flexible enough to incorporate different neural dynamics that have been observed at the cellular level in electrophysiological recordings.

The paper is accompanied by a software tool [11] that enables readers to visualize the effects of different model parameters on the neural dynamics. Many more neural dynamics can be simulated using the model and readers are encouraged to experiment with different network parameters. The paper and the tool illustrate how dynamical and spiking responses of neurons can be derived directly from a network objective or energy functional of continuous-values neural variables. The general approach offers an elegant way to design neuromorphic machine learning algorithms by bridging the gap that currently exists between bottom-up models that can simulate biologically realistic neural dynamics but do not have a network-level representation, and top-down machine learning models that start with a network loss function, but reduce the problem to the model of a non-spiking neuron with static nonlinearities.

In this regard, machine learning models are primarily developed with the objective of minimizing the error in inference by designing a loss function that captures dependencies among variables, for example, features and class labels. Learning in this case, as pointed out in [12], consists of adapting weights in order to associate low energies (losses) to observed configurations of variables, and high energies (losses) to unobserved ones. The non-differentiable nature of spiking dynamics makes it difficult to formulate loss functions involving neural variables. Neuromorphic algorithms currently work around this problem in different ways, including mapping deep neural nets to spiking networks through rate-based techniques [39, 40], formulating loss functions that penalize the difference between actual and desired spike-times [41, 42], or approximating the derivatives of spike signals through various means [43, 44, 45]. Formulating the spiking dynamics of the entire network using an energy function involving neural state variables across the network would enable us to directly use the energy function itself for learning weight parameters; and forms the basis for our future work. Since the proposed energy function encompasses all the neurons in the network, and not just the ‘visible neurons’ as in most neuromorphic machine learning algorithms, it can potentially enable easier and more effective training of hidden neurons in deep networks. Moreover, it would allow us to incorporate and experiment with biologically relevant neural dynamics that could have significant performance and energy benefits.

### 4.1 Relation with other neural networks and spiking neuron models

The network energy functional bears similarity with the Ising Hamiltonians used in Hopfield networks [46], Boltzmann machines [47] or spin-glass models [48], but contains an additional integral term ∫ Ψ(.)*dv* as in continuous-time Hopfield networks with graded neurons [49]. However, unlike in continuous-time Hopfield networks where Ψ^−1^(.) is assumed to be a saturation/squashing function of a rate-based representation, the role of Ψ(.) in the proposed model is to implement a barrier or a penalty, such that the neural responses can produce spiking dynamics. This enables us to obtain neural responses at the level of individual spikes instead of average rate-based responses; and allows for a more fine-grained control over the spiking responses of the network. The saturation (squashing) function, on the other hand, is implemented by the bound constraints on the Growth Transform updates, and hence the network is not limited to choosing a specific form of saturation non-linearity (e.g. sigmoid).

The energy-based formulation described in Section 2.1 could also admit other novel interpretations. For instance, for the form of Ψ(.) considered in 18, the barrier function can be rewritten as 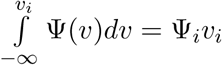,where Ψ_*i*_ = 0 *A* if *v*_*i*_ ≤ 0 *V* and Ψ_*i*_ = *I*_Ψ_ *A* if *v*_*i*_ > 0 *V*. For a continuous-time implementation (discrete-time step that is sufficiently small), *v*_*i*_(*t*) will be reset as soon as it reaches the threshold (0 *V*), and will not exceed the threshold. In this case, we can write Ψ_*i*_(*t*) *≥* 0 and Ψ_*i*_(*t*)*v*_*i*_(*t*) = 0 ∀*t*. This is equivalent to Karush-Kuhn-Tucker (KKT) conditions. Thus the spike events Ψ_*i*_(*t*), *i* = 1, …, *M*, act as the KKT multipliers corresponding to the *M* inequality constraints *v*_*i*_(*t*) *≤* 0, *i* = 1, …, *M* [50], *encoding the sensitivity of the i*-th neuron to the constraint.

Also, if we consider the spike response as a displacement current, we can write

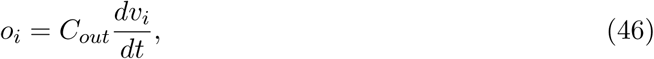

where *C*_*out*_ is the membrane capacitance. Note that *o*_*i*_ is the analog spike response current output and is different from Ψ(*v*_*i*_), which is the binary spike event. Then the membrane potential for the continuous-time Growth Transform neuron model in Table 3 can be rewritten as

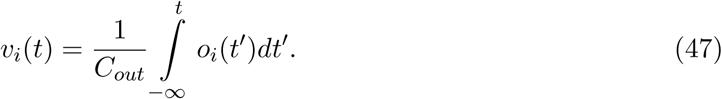

Thus according to this interpretation, the communication between neurons takes place using current waveforms, similar to integrate-and-fire models, and the current waveforms can be integrated at the post-synaptic neuron to recover the membrane potential. Note that the remapping between **W** and **Q** (described in Section 2.1) would still hold, since we are transmitting analog spike current waveforms, and not post-synaptic current waveforms such as exponentially decaying functions, *α*-functions or simplified current pulses (digital bits) used in integrate-and-fire models [51, 52].

### 4.2 Implication of remapping on neuromorphic architectures

In the proposed neuron model, we abstracted out the essential aspects of spike generation and transmission that can replicate neural dynamics, and remapped synaptic interactions to an energy-based framework. As a result of the remapping procedure, the coupling matrix **Q** in our proposed model is proportional to the inverse of the synaptic weight matrix **W**. This paves the way for developing novel neuromorphic learning algorithms in the **Q**-domain that involves sparse local analog connectivity, but which actually translates to fully-connected non-sparse global connectivity in the **W**-domain. Thus, adapting one synaptic connection in the **Q**-domain, in this case, will be equivalent to adapting multiple synapses in the **W**-domain. Learning in the **Q**-domain will be a topic for future research.

### 4.3 Benefits of decoupling neurodynamical parameters

A key advantage of the proposed framework is that it enables the decoupling of the three neurodynamical parameters - network solution, spike shapes and transient dynamics. Thus while the solution to the energy functional is determined by the coupling matrix **Q** and the stimulus vector **b**, independent control of the modulation function allows users to program the trajectory to the solution, which could be determined by an optimization process that is different from optimizing the energy functional. Some examples of these alternate objectives could be:

- A hybrid spiking network comprising neurons of different types (tonic spiking, bursting, non-spiking, etc.), as illustrated in Section 3.1. The network would still converge to the same solution, but the spiking dynamics across the network could be exploited to influence factors such as speed, energy efficiency and noise-sensitivity of information processing.
- Optimization of some auxiliary network parameter, e.g., the total spiking activity. A related example (although not optimized w.r.t. any objective function) was illustrated in Section 3.6 for a simple associative network. In this example, the network recalled the same set of patterns and classified MNIST images using two different time-evolutions of the modulation function corresponding to the presence and absence of global adaptation. In this case, it had the benefit of using fewer spikes to achieve better recall when a modified decoding metric was used.
- Effects of neurotransmitters and metabolic factors that have been known to affect the properties, activity and functional connectivity of populations of neurons. These factors endow the same network with the flexibility to generate different output patterns and produce different behaviors for the same stimulus [53, 54].
- Effects of diffusion processes or glial processes, that have been known to modulate response properties and synaptic transmission in neurons, influencing information processing and learning in the brain [55, 56].

## Conflict of Interest Statement

The authors declare that the research was conducted in the absence of any commercial or financial relationships that could be construed as a potential conflict of interest.

## Author Contributions

AG and SC contributed to the conception and design of the study; AG and DM conducted the simulations; AG, DM and SC designed the MATLAB interface; AG and SC wrote the first draft of the manuscript. All authors contributed to the manuscript revision, read and approved the submitted version.

## Funding

This work was supported in part by a research grant from the National Science Foundation (ECCS: 1935073). D.M. was supported by a research grant from the National Institutes of Health (5R21EY02836202).

## Acknowledgments

The authors would like to thank Dr. Kenji Aono at the Electrical and Systems Engineering department, Washington University, for developing a GPU version of the GT neural network model which is also included with the accompanying software toolbox [11].

### Appendix A

We can rewrite (1) for a sequence (*v*_*i,n*−*N*+1_, *v*_*i,n*−*N*+2_, …, *v*_*i,n*_) of size *N* to obtain

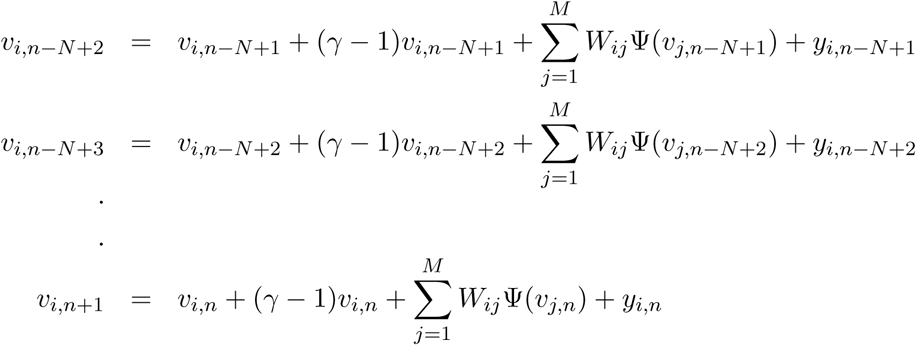

Summing over the time-steps and dividing by the total number of time-steps, we get

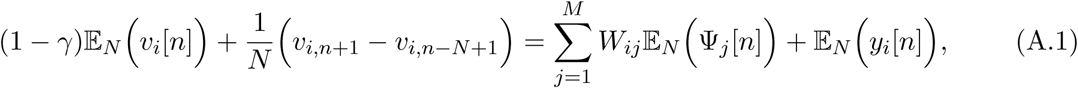

where 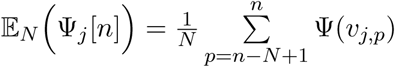. Since the neural responses are assumed to be bounded at all times, as *N* → *∞*, the second term in (A.1) approaches zero, so that we can rewrite (A.1) as

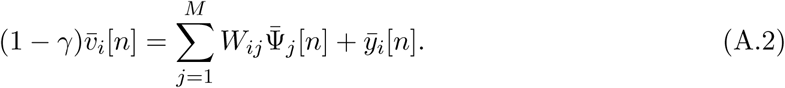

### Appendix B

#### Proof of Proposition I

We can decompose the scalar variable *v*_*i,n*_ as 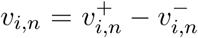 where 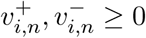. The following additional constraint

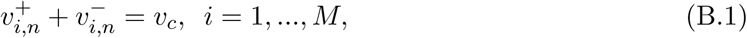

imposed on the variables 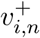 and 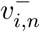 would then always ensure that (10) is satisfied. Then we can write

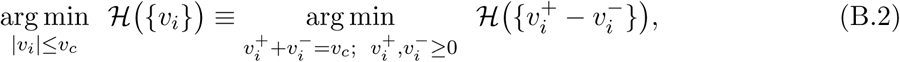

We can solve the equivalent optimization problem under the linear constraint specified by (B.1) and the non-negativity constraints, using an iterative multiplicative update called Growth Transforms - a fixed point algorithm for optimizing a Lipschitz continuous objective function under similar equality constraints [57, 20].

#### Baum-Eagon Growth Transformations

Baum-Eagon Growth Transforms [20] are a class of fixed-point algorithms for iteratively optimizing a Lipschitz continuous objective function *ℋ* (*{v*_*ik*_*}*) that is constrained over a domain *D* defined by

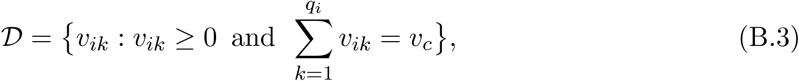

where *q*_1_, *q*_2_, …, *q*_*M*_ is a set of non-negative integers and *M* is a positive integer which denotes the number of linear constraints in *D*. For a Lipschitz continuous cost function *ℋ* (*{v*_*ik*_*}*) where *v*_*ik*_ *∈ 𝒟, i* = 1, …, *M*, there exists a growth transformation *σ*(.) such that

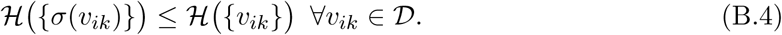

This transformation takes the following form

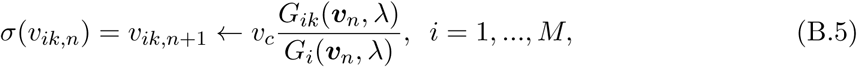

Where

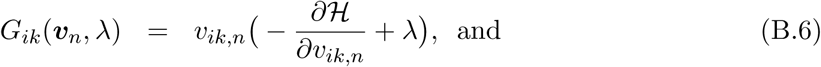

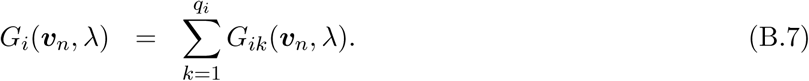

An admissible value for the constant *λ* is such that for any *v*_*ik,n*_ *∈ 𝒟, G*_*ik*_(***v***_*n*_, *λ*) *≥* 0 and *G*_*i*_(***v***_*n*_, *λ*) *>* 0 [57].

#### Growth Transform neuron model updates

Assuming that the partial derivatives for the network energy functional in (8) are bounded, the optimization problem for the proposed model is equivalent to the one in (B.2), being constrained on a domain equivalent to *𝒟*, where *q*_1_ = *q*_2_ = … = *q*_*M*_ = 2, and 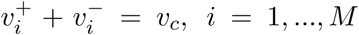 are the *M* linear constraints. Considering 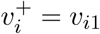 and 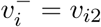, we can rewrite the update equations in (B.5) in terms of the new optimization variables corresponding to (B.2) to obtain the following discrete-time updates

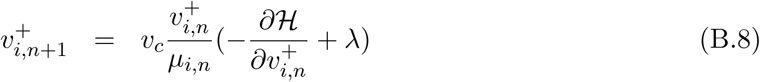

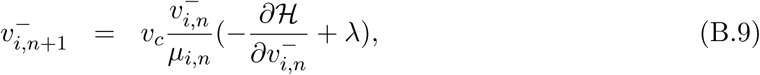

Where

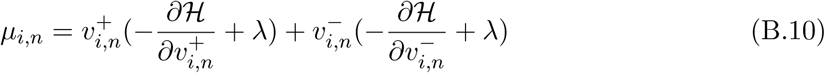

is a normalization factor that ensures 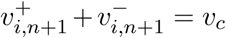. Here, *λ* (which has the unit of current) is admissible iff 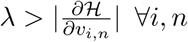, which will ensure 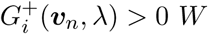 and 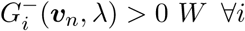. From (B.8) and (B.9), using 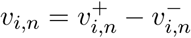 and (B.1), we can easily show

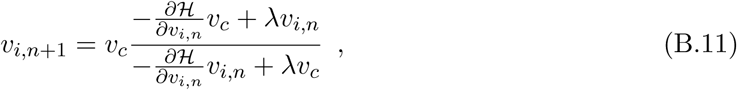

where we have used the relation 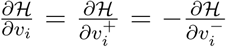. This dynamical system model, derived from the Growth Transform updates outlined in (B.8) and (B.9), ensures that (11) holds, with equality being satisfied iff *v*_*i,n*_ is a critical point of *H*.

Rearranging the terms in (B.11), we get

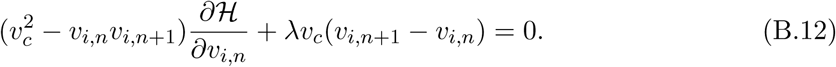

Rewriting B.12 for a sequence of time-indices *p* = *n* − *N* + 1, *n* − *N* + 2, …, *n*, of size *N*, we get

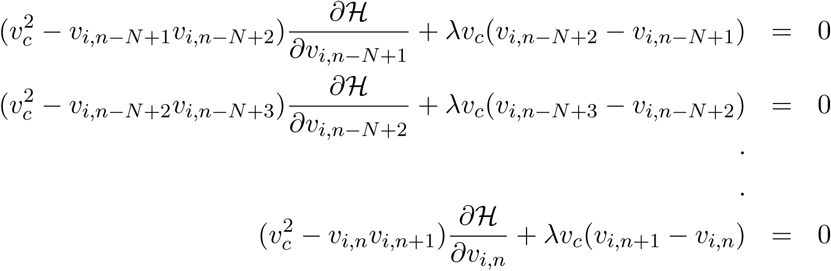

Summing over the time-steps and dividing by the total number of time-steps, we get

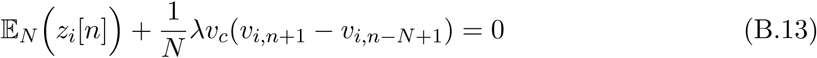

where 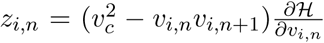. As *N* → *∞*, since *v*_*i,n*_ are bounded ∀*i, n*, we have for the *n*-th time-window

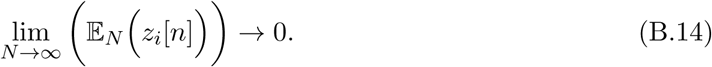

### Appendix C

#### Asymptotic encoding for non-saturating GT neurons

For neurons with responses *v*_*i,n*_ *>* −*v*_*c*_ ∀*n* (which includes spiking neurons as well as non-spiking neurons that do not cross the threshold), we define 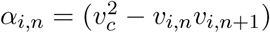. Since |*v*_*i,n*_| *< v*_*c*_ ∀*n, α*_*i,n*_ *>* 0 ∀*n*. We can sum the criterion (B.12) for time-steps *p* = *n* − *N* + 1, *n* − *N* + 2, …, *n*, to write

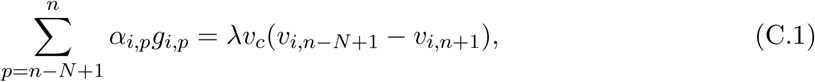

where 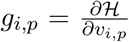. For *p* = *n* − *N* + 1, *n* − *N* + 2, …, *n*, we can decompose the instantaneous gradient term *g*_*i,p*_ as follows

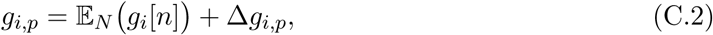

where Δ*g*_*i,p*_ is a zero-mean sequence such that 𝔼_*N*_ (Δ*g*_*i*_[*n*]) = 0. Then combining (C.1) and (C.2),

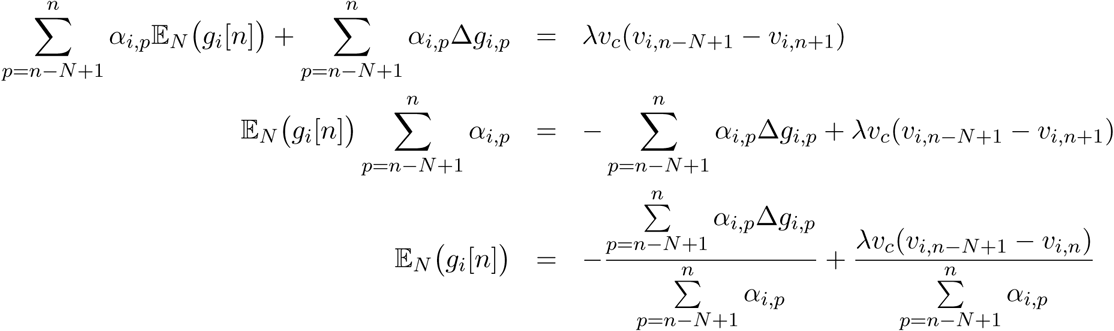

Since 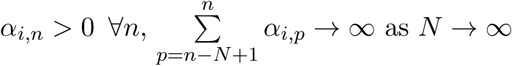. Also, |*v*_*i,n*_| *< v*_*c*_ ∀*n* leads to

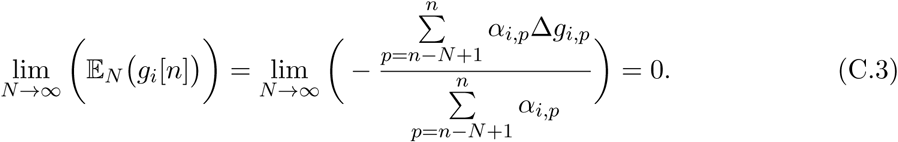

The last result uses the non-pathological case that due to the bounded property of Δ*g*_*i,p*_, the sequence *α*_*i,p*_Δ*g*_*i,p*_ does not grow as fast as the denominator sequence *α*_*i,p*_.

### Appendix D

#### Continuous-time Growth Transform dynamical system

In order to derive the complete dynamical system model for the Growth Transform neuron, we apply a useful property of Growth Transforms. The Growth Transform mapping *σ*(.) homotopically minimizes the value of the cost function *ℋ* [9] as shown below

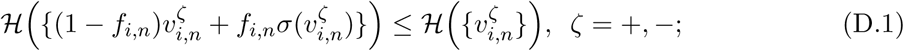

where 0 *< f*_*i,n*_ *≤* 1. This leads to the updated discrete-time equations for the new optimization variables for minimizing *ℋ* (.)

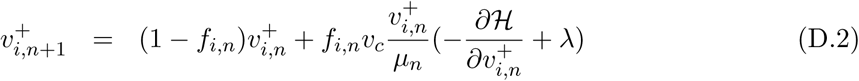

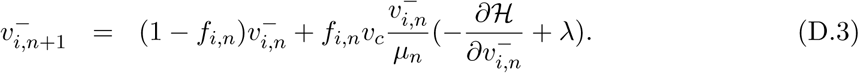

From (B.11), (D.2) and (D.3), we have

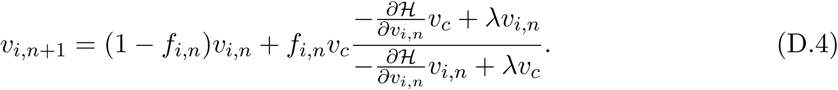

We define *τ*_*i,n*_ = Δ*t*(1*/f*_*i,n*_ −1), where Δ*t* is the time-increment in seconds between two time-steps as defined previously in (1). Then since 0 *< f*_*i,n*_ *≤* 1, we have *τ*_*i,n*_ ∈ [0, *∞*) s. *τ*_*i,n*_ can be considered to be the time-constant for the *i*-th neuron at the *n*-th time-step, and the discrete-time dynamical systems model in (D.4) can be written as

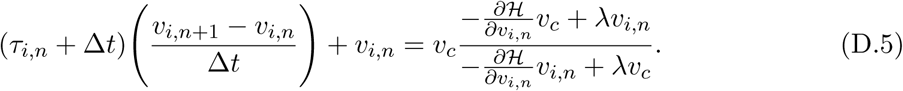

Since the *n*-th time-step corresponds to time *t* = *n*Δ*t, v*_*i,n*_ ≡ *v*_*i*_(*n*Δ*t*) and (D.5) can be rewritten as

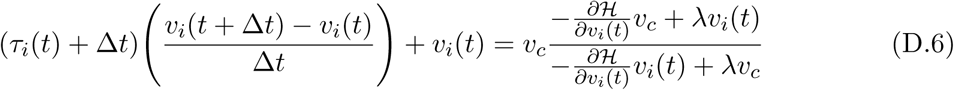

In the limiting case when Δ*t* → 0 s, this reduces to the following continuous-time dynamical system model [20]

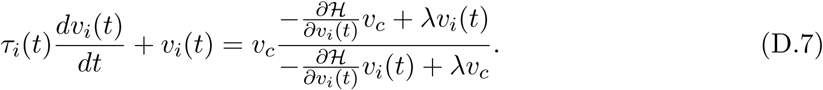

## Notes

https://github.com/aimlab-wustl/growth-transform-NN

